# Macrophage-expressed Neuropilin 2 promotes plaque formation in ApoE knockout mice and is essential for pro-inflammatory macrophage polarisation and gene expression

**DOI:** 10.1101/2023.09.27.558806

**Authors:** Jacob Fernando-Sayers, Jennifer L. Harman, Matthew C. Gage, Ian C. Zachary, Caroline Pellet-Many

**Affiliations:** Centre for Cardiovascular Biology, University College London, 5 University Street, London WC1E 6JF, UK; Department of Comparative Biomedical Sciences, Royal Veterinary College, 4 Royal College Street, London NW1 0TU, UK

**Keywords:** Neuropilin 2, macrophage polarisation, macrophage migration, atherosclerosis, transcriptomics

## Abstract

**Aims:** Atherosclerosis is a chronic inflammatory disease causing the narrowing of arteries, leading to ischaemic heart disease. It is characterised by the subendothelial retention, and modification, of lipoproteins by macrophages, highly plastic cells which undergo polarisation to a pro-inflammatory phenotype in response to cytokines and other environmental stimuli.

Neuropilin-2 (NRP2) is a cell-surface co-receptor with essential roles in angiogenesis and axonal guidance, that is also expressed by macrophages. However, the role of NRP2 in macrophage function in the development of atherosclerosis has not been studied.

**Methods and results:** The role of NRP2 in modulating macrophage polarisation and signalling was assessed using functional assays and transcriptome analysis in macrophages obtained from mice with macrophage-specific deletion of *Nrp2* (*Nrp2-KO^Mac,EYFP^*). These mice were further crossed with pro-atherogenic Apolipoprotein E-deficient mice to produce *Nrp2-KO^Mac,Apoe-/-,EYFP^* mice, which were fed a high-fat diet (HFD) for 16 weeks. Plaque formation and composition were characterised using chemical and immuno-staining.

NRP2 was significantly upregulated upon differentiation of bone marrow progenitors into bone marrow derived macrophages (BMDM), and further upregulated by pro-inflammatory polarisation. Transcriptome analysis revealed that inflammatory signalling pathway genes, and genes regulating monocyte chemotaxis, were downregulated in *Nrp2-KO^Mac,EYFP^* BMDMs. HFD-induced plaque development was significantly reduced in *Nrp2-KO^Mac,Apoe-/-,EYFP^* mice. Additionally, plaques from those mice displayed features consistent with increased plaque stability, including reduced necrotic core area, plaque lipid content and increased cap thickness.

**Conclusions:** Macrophage-derived NRP2 is proatherogenic, likely resulting from its ability to positively regulate pro-inflammatory pathways and macrophage migration. Targeting NRP2 expressed on the surface of macrophages could therefore offer a novel therapeutic approach for reducing the disease burden associated with atherosclerosis.

## 1. Introduction

Cardiovascular disease (CVD) is the leading cause of death worldwide and is expected to be a globally escalating problem for many years ^1, 2^. CVD is predominantly the clinical manifestation of atherosclerosis, a chronic and progressive inflammatory disease of large to medium sized arteries ^3^, characterised by the narrowing of blood vessels due to a build-up of plaque comprising lipids, extracellular matrix (ECM) and several cell types such as endothelial cells ^4^, smooth muscle cells ^5, 6^ and leukocytes ^7^.

Macrophages are highly heterogenous and plastic mononuclear phagocytes derived from monocytes, with a primary role in innate immunity, and additionally important in tissue homeostasis, regeneration and remodelling in physiological and pathophysiological settings ^8^. Both tissue-resident and monocyte-derived macrophages are involved during all stages of atherosclerotic plaque formation ^9^. Following monocyte migration into the arterial wall sub-endothelium, monocytes differentiate into macrophages in response to locally secreted cytokines ^9^. There, they drive plaque progression via local proliferation ^10, 11^, the uptake, accumulation and oxidation of cholesterol, and positive feedback to both enhance monocyte recruitment through secretion of chemokines such as Chemokine (C-C motif) ligand 5 (CCL5) and chemokine (C-X-C motif) ligand 1 (CXCL1) and expression of the chemokine receptors, chemokine (C-X3-C motif) receptor 1 (CX3CR1) and C-C chemokine receptor 2 (CCR2) ^12, 13^, thus committing to a pro-inflammatory phenotype ^14^. Improving our understanding of the mechanisms involved in the regulation of macrophage phenotype is therefore a prerequisite for identifying novel interventions to prevent atherogenesis.

Neuropilins, NRP1 and NRP2, are structurally related transmembrane receptors with well-established roles in blood vessel formation, lymphangiogenesis and the formation and homing of nerve fibres ^15-17^. More recent evidence indicates an important role for NRPs in regulating the function of immune cells, including macrophages ^18^. Genetic *Nrp1* depletion in mouse tumour-associated macrophages (TAMs) arrests their pro-angiogenic and immunosuppressive functions thus decreasing tumour growth and metastasis in Lewis lung carcinomas (LLC) ^19^ and glioma models ^20^. Furthermore, myeloid-specific deletion of *Nrp1* enhanced inflammatory cytokine production via Toll-like receptor 4 (TLR4)-Nuclear factor kappa B (NF-κB) pathways resulting in increased sepsis following intraperitoneal injection of lipopolysaccharides (LPS) ^21^. Additionally, the same group also found that loss of *Nrp1* in macrophages increased insulin resistance via NOD-, LRR- and pyrin domain-containing protein 3 (Nlrp3) inflammasome priming and activation ^22^. While these findings identify a role for NRP1 in suppressing pro-inflammatory macrophage functions, the regulation and role of NRP2 in macrophages remain less understood. NRP2 expression is undetectable in monocytes but is significantly upregulated as they differentiate into macrophages ^23^. Roy et al. demonstrated that *Nrp2* myeloid-specific deletion impairs efferocytosis of TAMs and tumour growth as well as decreasing the expression of immunosuppressive and tumour-promoting genes ^23^, thus suggesting immunosuppressive functions for NRP2 in TAMs. However, studies of *Nrp2* regulation in macrophages have yielded contrasting results: LPS stimulation of airway macrophages upregulated *Nrp2* in a mouse model ^24^, whereas LPS downregulated macrophage *Nrp2* expression *in* human macrophages ^25^.

To investigate the role of NRP2 in atherogenesis, we initially characterised expression of Neuropilin 2 gene and protein during the differentiation of monocytes into macrophages and following macrophage polarisation ^26^. We next generated a mouse model with myeloid-specific knockout (KO) of *Nrp2* (*Nrp2-KO^Mac,EYFP^*) and crossed to the pro-atherogenic Apolipoprotein E (Apo-E) knockout mice model (*Nrp2-KO^Mac,ApoE-/-,EYFP^*) to investigate the impact of *Nrp2* loss in macrophages on atherosclerotic plaque development. Additionally, we performed transcriptome analysis of *Nrp2-KO^Mac,EYFP^* BMDMs to identify differentially expressed genes (DEGs) and pathways modulated by *Nrp2*.

## 2. Methods

### 2.1 Generation of Mouse models

All procedures were performed in accordance with the UK Animals (Scientific Procedures) Act 1986 by the Personal License holder under the Home Office Project License PB38E126B and NC3Rs recommendations. Mice were bred and maintained in a pathogen-free biological services unit animal facility (UCL Central Biological Services Unit) with a 12-hour light-dark cycle, and ad libitum access to food and water. The C57BL/6J mouse line expressing Cre-recombinase under the control of the Lysozyme M promoter (LysM-Cre) was crossed to the C57BL/6J *Nrp2^flox^* mouse, both imported from Jackson Laboratory (#004781 and #006697, respectively). Those were then crossed to the *R26R-EYFP*^+/+^, *ApoE*^−/−^ mouse line which was a generous gift from the lab of Prof. Gary Owens’ Laboratory (Virginia, USA) and resulted from the cross of mouse lines #006148 (R26R-EYFP) and #002052 (*ApoE* KO) respectively (Jackson Laboratory). We generated myeloid-specific deletion of *Nrp2* and EYFP expression (to enable lineage tracing) on a Wild-Type (WT) or proatherogenic background *Nrp2-KO^Mac,EYFP^* and *Nrp2-KO^Mac,ApoE-/-,EYFP^* respectively. Littermates expressing LysM-Cre and EYFP, but WT NRP2, were used as controls (*WT^Mac,EYFP^*), with similar controls on a proatherogenic background (*WT^Mac,ApoE-/-,EYFP^*). Genotyping primers sequences are specified in Supplementary Table I.

### 2.2 High-fat Diet protocol

8-week old *Nrp2-KO^Mac,ApoE-/-,EYFP^* mice were weighed and then fed, ad libitum, a high-fat diet (HFD) (21.4% fat, 0.15% cholesterol; Special Diet Services, 823119) for 16 weeks. At 24 weeks old, these mice were weighed and euthanised via carbon dioxide (CO_2_) overdose followed by cervical dislocation.

### 2.3 L929-conditioned medium (LCM) preparation

Macrophage colony-stimulating factor-transduced L929 cells (obtained from the Pineda-Torra Lab, Division of Medicine, UCL) were cultured in T-175 cm^2^ tissue culture flasks (Greiner Bio-One, 660175) with Dulbecco’s Modified Eagle Medium (DMEM) (Gibco, 41966-029) supplemented with 10% Foetal Bovine Serum (FBS) (Gibco, 10270-106) and 1% Penicillin (10,000 U/mL)/Streptomycin (10 mg/mL) (Gibco, 15140-122) in a humidified tissue culture incubator (37°C, 5% CO_2_). Medium was replaced every 2 days until the cells reached 90% confluence. Once the cells reached confluence, the medium (40 mL) was collected and replaced approximately every 10 hours over a 48-hour period. After each collection, the medium was centrifuged at 1000 x *g* for 5 minutes, to remove cells and debris, and stored at 4°C until all collections had been made. After the final collection, all collections were pooled (200 mL per T-175 cm^2^ tissue culture flask) and filtered through a 0.45 μm vacuum filter unit (ThermoFisher Scientific, 1670045) to produce the L929-conditioned medium (LCM), which is high in M-CSF and used experimentally to differentiate bone marrow progenitors into macrophages. The LCM was then stored at -20°C until required for experiments.

### 2.4 Macrophage culture, differentiation and polarisation

Bone-marrow was extracted from the tibiae and femurs of 10- to 12-week-old *Nrp2-KO^Mac,EYFP^* and control WT sibling mice as previously described ^26^. The tibiae and femurs were stripped of adherent tissue, epiphyses were removed, and bone marrow extracted via flushing with DMEM GlutaMax™ (Gibco, 10566-016). Medium containing bone marrow was centrifuged for 5 minutes at 310 x *g* and the supernatant was discarded. After pelleting the bone marrow, red blood cells were eliminated by resuspending the pellet in Red Blood Cell Lysis Buffer (Sigma-Aldrich, R7757) at RT for 5 minutes. The remaining cells were then centrifuged for 5 minutes at 310 x *g* and resuspended in bone marrow-derived macrophage (BMDM) differentiation medium, consisting of DMEM GlutaMax™ supplemented with 20% FBS (Gibco, 10270-106), 30% LCM, and 20 µg/mL Gentamycin (Gibco, 10131-035). Cells were counted and seeded at a density of 1 x 10^6^ cells or 5 x 10^6^ cells per well of a 6-well plate. After 3 days in culture, cells were supplemented with additional BMDM differentiation medium and after 5 days of culture the cells were washed twice with sterile PBS before adding fresh BMDM differentiation medium. After 7 days of culture, macrophages are fully differentiated with >90% of cells expressing the cell surface marker CD11b, and 99.5% positive for the macrophage marker, F4/80, and described as proliferative non-activated (M0) macrophages ^26^. At this stage, some cells were harvested to assess the expression of the EYFP marker under the control of the constitutive LysM-Cre recombinase, we found that all cells were positive for EYFP. Following macrophage differentiation, BMDM differentiation medium was replaced with BMDM activation medium consisting of DMEM GlutaMax™ supplemented with 10% FBS and 20 μg/mL Gentamycin. M0 macrophages were then polarised using 20 ng/mL IFNγ (Peprotech, 315-05) and 100 ng/mL LPS, or 20 ng/mL IL4 (Peprotech, 214-24) to produce pro-inflammatory (M1) or pro-angiogenic, immunosuppressive (M2) macrophages respectively. Macrophages were left to polarise for 24 hours prior to RNA analysis, or 48 hours for protein analysis.

### 2.5 Real-time (RT) quantitative PCR (qPCR)

BMDMs in tissue-culture plates were placed on ice, washed twice with ice-cold PBS. Prior to RNA extraction, BMDMs were disrupted via scraping in RLT buffer and homogenised through QIAshredder spin columns (Qiagen, 1011711). Total RNA was then extracted from BMDMs using the RNeasy Mini Kit (Qiagen, 74106) as per the manufacturer’s instructions. The concentration and purity of the RNA samples were analysed using a NanoDrop spectrophotometer (ThermoFisher Scientific) and 250 ng of RNA from each sample was used for subsequent complementary DNA (cDNA) synthesis. Following treatment with DNase, cDNA synthesis was performed using the QuantiTec® Reverse Transcription Kit (Qiagen, 205311) as per the manufacturer’s instructions. Samples were then diluted (1:10) in ddH_2_O and stored at -20⁰C.

Genes of interest were amplified and analysed via qPCR using Brilliant III SYBR® Green qPCR master mix (Agilent, 600882) on the MX3000p qPCR detection system (Stratagene, Agilent Technologies). Primers (Supplementary Table II) were purchased from ThermoFisher Scientific and were designed to span or flank introns and anneal at 60⁰C. All reactions had an initial denaturation step at 95⁰C for 3 minutes followed by 40 cycles of denaturation at 95⁰C for 10 seconds, combined annealing and extension at 59-61⁰C for 15 seconds. Product specificity was determined by visualisation of the amplified products using agarose gel electrophoresis and by melting curve analysis (Tm). Expressions of genes of interest were normalised to the reference genes, *Actb* or *Gapdh*. Relative gene expression was calculated using the double delta cycle threshold (ddCt) method of quantification as described previously ^27^.

### 2.6 Western blot

Following polarisation, protein lysates were extracted from BMDMs using Radioimmunoprecipitation assay (RIPA) buffer (Sigma-Aldrich, R0278) complimented with protease inhibitor (Roche Diagnostics, 04693116001), phosphatase inhibitors (Sigma-Aldrich, P5726 and P2850) and reducing agent tris(2-carboxyethyl)phosphine (TCEP) (Sigma-Aldrich, 646547). Lysates were centrifuged at full speed for 10 minutes at 4⁰C and the total protein concentration in the supernatant was quantified using the detergent compatible (DC) protein assay (BioRad, 500-0114). Equal amounts of protein were mixed with anionic detergent (4x LDS loading buffer, Novex, NP0007) and samples were heated to 95⁰C for 3 minutes for protein denaturation. Samples were run on 4-12% Bis-Tris polyacrylamide gels (Novex, NP0322) in NuPage® SDS MOPS running buffer (ThermoFisher Scientific, NP0001). Proteins were then electro-transferred onto polyvinylidene fluoride (PVDF) membranes (ThermoFisher Scientific, LC2002). The protein-bound PVDF membranes were blocked with blocking solution (5% (w/v) non-fat dried milk in 1X PBS Tween™-20 (PBST) (ThermoFisher Scientific, 28352)) for 1 hour at room temperature (RT) on a rocking platform and then incubated with the primary antibodies (Supplementary Table III) overnight at 4⁰C. Primary antibodies were diluted according to manufacturer’s recommendations in blocking solution. Following overnight incubation, the membranes were washed in PBST and subsequently incubated with horseradish peroxidase (HRP)-conjugated secondary antibodies (Supplementary Table III) at RT for 1 hour on a rocking platform. All secondary antibodies were diluted 1:5000 in blocking solution. Membranes were washed four times for 5 minutes in PBST and incubated with Immobilon® Forte Western HRP Substrate (Merck, WBLUF0100) for 3 minutes. Membranes were then placed in an autoradiography cassette and exposed to autoradiography film (Merck, GE28-9068-35) in a dark room. Developed films were scanned and densitometry analysis was performed using the FIJI analysis software (v.2.1.0/1.53c). Bands of protein of interest were normalised against appropriate loading control proteins, and expression of protein of interest is displayed as a ratio relative to expression of the loading control.

### 2.7 Fluorescent immunocytochemistry (ICC) of cultured BMDM

BMDM were differentiated and polarised on glass cover slips in 6-well plates. Following the treatment, cover slips were washed with ice cold PBS and BMDMs were fixed in 4% (w/v) paraformaldehyde (PFA) (in PBS) (Sigma-Aldrich, P6148) for 15 minutes at RT. After a brief wash in PBS, cells were permeabilised with 0.1% TritonX100 (ThermoFisher Scientific, 85111) (in PBS) for 10 minutes and then incubated in blocking solution (1% BSA in PBST with 10% either donkey or goat serum depending on the species in which the secondary antibody was raised) for 1 hour at RT. Coverslips were then incubated overnight at 4⁰C with primary antibodies (Supplementary Table III) diluted (1:100) in blocking solution. Following overnight incubation, samples were washed 5 times in PBS and incubated with fluorescent secondary antibodies diluted (1:500) in blocking solution in the dark for 1 hour at RT. Finally, coverslips were rinsed with PBS and mounted onto glass slides using ProLong® Diamond Antifade reagent with DAPI mounting medium (ThermoFisher Scientific, P36962). Images were generated using the Leica TCS SP8 confocal microscope system or an upright fluorescence microscope and analysed using the FIJI analysis software (v.2.1.0/1.53c).

### 2.8 Chemotactic Migration assay

BMDM were differentiated as described above and adapted from ^28^. After 7 days of culture, BMDMs were washed twice in warmed PBS, gently detached from the culture dish using a cell scraper, transferred to a 15mL tube and centrifuged at 310 x *g* for 5 minutes at RT. The cell pellet was then resuspended in DMEM GlutaMax™ supplemented with 10% FBS and 20 μg/mL gentamycin and counted using an ADAM-MC™ Automatic Cell Counter (DigitalBio, ADAM-MC). After determining cell concentration, additional medium was added to reach a final concentration of 1x10^6^ cells/mL.

600 μL of DMEM GlutaMax™ supplemented with 10% FBS, 20 μg/mL gentamycin, and either 100 ng/mL recombinant mouse Monocyte chemoattractant protein-1 (MCP-1, CCL2) (Bio-Rad, PMP35) or vehicle control (ddH_2_O), was added into the bottom of the wells of a 24-well plate. Polyethylene terephthalate (PET) transwell inserts with 8 μm pores (Corning, 353097) were inserted into each well and 500 μL of BMDM cell suspension (5x10^5^ cells) was added into each transwell and left to migrate for 20 hours in a humidified tissue culture incubator at 37⁰C, 5% CO_2_. After 20 hours, the transwells were removed and non-migrated cells wiped away using a cotton bud by gentle rotation. The transwell membranes were fixed and stained using the REASTAIN® Quick-Diff Kit (Reagena, 102164), in accordance with the manufacturers’ instructions. After staining, the transwells were dried at RT for 24 hours, before imaging with a brightfield microscope at 80X magnification. The number of migrated BMDMs was counted manually, using the cell counter plugin on the FIJI analysis software in 5 microscopic fields per membrane. Results are displayed as the average number of migrated BMDMs across the 5 microscopic fields, avoiding the edge of the inserts.

### 2.9 Aorta removal and en-face Oil-Red-O staining

Following euthanisation, and tissue processing as described above, unwanted abdominal organs, the diaphragm and the lungs were removed under a dissecting stereomicroscope to fully expose the heart and entire aorta. The heart and aorta were extracted, washed in PBS, and separated by cutting the ascending aorta at the base of the heart. The heart was then placed in 4% (w/v) PFA on a rotating platform for 24 hours at 4⁰C before embedding for histological analysis (as described in the paragraph below). The entire aorta was dissected free of periaortic adipose tissue, and the adventitia carefully peeled away from the media. The aorta was then fixed in 4% (w/v) PFA (in PBS) at 4⁰C overnight. In preparation for Oil-Red-O staining, a longitudinal incision was made along the lesser curvature of the aortic arch that extended down to the bifurcation of the aorta at the common iliac arteries.

Prior to Oil-Red-O staining, aortas were washed in ddH_2_O for 1 minute to remove excess PFA and incubated in 60% isopropanol for 2 minutes to sustain neutral lipids and facilitate staining. Aortas were then stained with filtered 60% Oil-Red-O (in ddH2O) in the dark for 12 minutes at RT on a rotating platform. After staining, the aorta was briefly rinsed in 60% isopropanol to remove excess Oil-Red-O and stored in ddH_2_O in preparation for imaging. Aortas were mounted onto glass slides and imaged under a stereomicroscope. Images were stitched together and analysed using the FIJI analysis software. Lesion areas were manually quantified as areas of Oil-Red-O staining and recorded as a percentage of the total area of the aorta.

### 2.10 Heart processing and sectioning for root staining

Following 24-hour fixation in 4% (w/v) PFA (in PBS), hearts were washed briefly in PBS incubated in 30% sucrose (Sigma-Aldrich, S0389) in PBS for 24 hours at 4⁰C on a rotating platform. Hearts were then incubated in a 50:50 solution of 30% sucrose:Optimal cutting temperature (O.C.T.) compound (VWR, 361603E) for 1 hour at RT on a rotating platform, followed by a final incubation in O.C.T. for 1 hour at RT. The apex of the heart was cut to allow the heart to sit upright with the aortic root cross-section sitting in the transverse plane and placed in plastic moulds before being embedded in O.C.T. on dry ice. Embedded hearts were stored at -80⁰C until cryosectioning. 12 μm thick transversal sections were obtained using a cryostat, in the base-apex direction. Once it was confirmed that the aortic root had been reached, using a light microscope, serial sections were mounted onto SuperFrost Plus Adhesion microscope slides (ThermoFisher Scientific, 10149870), placed on dry ice and stored at -80⁰C until needed for staining.

### 2.11 Haematoxylin and eosin staining of aortic root cryosections

To assess cap thickness and necrotic core area within atherosclerotic plaques, aortic root sections were stained with haematoxylin and eosin (H&E). Aortic root cryosections were dried at RT for 10 minutes and fixed in acetone (Acros organics, 423245000) at -20⁰C for 3 minutes, followed by 80% methanol at 4⁰C for 5 minutes. Slides were then washed in PBS at RT for 10 minutes. Aortic roots were stained in pre-filtered haematoxylin (Sigma-Aldrich, HHS32) for 1 minute, washed under running water for 5 minutes and finally stained in eosin (Sigma-Aldrich, HT110280) for 30 seconds. Aortic root sections were then dehydrated in an increasing ethanol series (5 minutes in each solution, from 70%, 80%, 90% and 100% ethanol) followed by incubation in xylene, and mounted with DPX mounting medium (Sigma-Aldrich, 06522) and glass cover slips. Aortic roots were imaged using an automated slide scanner (NanoZoomer, Hamamatsu), and cap thickness and necrotic core area were calculated using the NDP.view2 Viewing Software (Hamamatsu, U12388-01). Cap thickness was calculated as the average cap thickness from at least 4 independent cap measurements taken from a minimum of 2 aortic root sections per mouse. Necrotic core area was calculated as the average percentage necrotic core coverage (relative to total plaque area) taken from at least 2 aortic root sections per mouse.

### 2.12 Oil-red-O staining of aortic root cryosections

To assess atherosclerotic plaque size and lipid content, aortic root sections were stained with Oil-Red-O. Aortic root cryosections were dried at RT for 10 minutes and washed in ddH_2_O for 1 minute to remove excess O.C.T. Slides were then placed in 60% isopropanol for 2 minutes to facilitate staining of neutral lipids. The aortic root sections were stained with filtered 60% Oil-Red-O (in ddH2O) in the dark for 12 minutes at RT. Sections were briefly rinsed in 60% isopropanol to remove excess Oil-Red-O and then further washed briefly in ddH2O to remove the isopropanol. Sections were counterstained with pre-filtered haematoxylin for 1 minute and washed under running water for 5 minutes. Aortic root sections were mounted with 90% glycerol (in PBS) and glass cover slips, and imaged under a stereomicroscope (Leica, MZ10F). Images were then analysed using the FIJI analysis software, where lipid content was manually quantified as percentage lipid relative to total root cross-sectional area, and reported as the average percentage lipid across 2-3 aortic root sections per mouse.

### 2.13 Fluorescent immunohistochemistry (IHC) of aortic root cryosections

To assess the cellular content of atherosclerotic plaques, in particular macrophage and SMC content, immunofluorescence staining was performed on aortic root sections. Aortic root cryosections were dried at RT and washed in ddH_2_O for 1 minute to remove excess O.C.T. Sections were then fixed in 4% (w/v) PFA (in PBS) for 15 minutes at RT, washed briefly in PBS and permeabilised in 0.1% Triton X100 (in PBS) for 10 minutes at RT. All primary antibodies used were raised in mice, therefore sections were blocked using a M.O.M.® (mouse-on-mouse) Immunodetection Kit (Vector Laboratories, BMK-2202), as per the manufacturers protocol, to block mouse immunoglobulins and reduce background fluorescence. Sections were incubated in M.O.M, Mouse IgG Blocking Reagent (Vector Laboratories, BMK-2202) for 1 hour at RT, washed twice for 2 minutes in PBS and incubated in M.O.M. Diluent (Vector Laboratories, BMK-2202) for 5 minutes at RT. Next, anti-GFP primary antibody (Supplementary Table III) was diluted (1:100) in M.O.M. Diluent and added to the sections, for overnight incubation at 4⁰C. Following overnight incubation, sections were washed 5 times in PBS and incubated with secondary antibody (Supplementary Table III), diluted (1:500) in M.O.M. Diluent, for 1 hour in the dark at RT. Excess secondary antibody was removed by washing twice in PBS for 2 minutes. Sections were incubated in fluorophore-conjugated alpha smooth muscle actin (αSMA) antibody (Supplementary Table III), diluted (1:100) in M.O.M. Diluent, for 3 hours in the dark at RT. Sections were washed twice in PBS for 2 minutes and mounted under glass cover slips using ProLong® Diamond Antifade reagent with DAPI mounting medium. Aortic roots were imaged using a fluorescent microscope.

### 2.14 RNA Sequencing

Bone marrow was extracted from the tibiae and femurs of three 10- to 12-week-old mice per genotype (wild-type and *Nrp2-KO^Mac, EYFP^*), pooled, and differentiated as described previously. On day 7 of culture, tissue culture plates containing M0 BMDMs were placed on ice, medium was removed, cells were washed twice with ice cold PBS and then disrupted through scraping in RLT buffer. Total RNA extraction and sample purity analysis were then performed using the protocol previously described. RNA samples were then transported, on dry ice, to UCL Genomics (London, UK) for RNA integrity analysis, library preparation (Kappa mRNA Hyper, Kappa Biosystems), sequencing (NovaSeq 6000 SP, Illumina) and pipeline analysis (SARTools ^29^).

Sequencing and computational analysis were performed by UCL Genomics. Libraries were sequenced, using an Illumina NovaSeq 6000 machine, and reads were de-multiplexed using Illumina bcl2fastq (Illumina). De-multiplexed reads were then aligned to the mouse GRCm38/mm10 sequence using STAR-2.7.9a, and filtered to remove singletons and unaligned reads. Analysis/identification of differentially expressed genes (DEGs) was performed with SARTools (a DESeq2 and EdgeR package). To correct for multiple comparisons, and reduce the probability of false positives, SARTools automatically sets an adjusted p-value threshold of 0.05. This analysis produced an individual up- and downregulated DEG list, which comprised a list of genes that were up- or downregulated in *Nrp2*-KO macrophages relative to expression in wild-type murine macrophages. Prior to conducting downstream analysis on the DEG lists, a tighter significance threshold and a log fold-change (LFC) cut-off (-log10(Padj) >2 and -0.5> LFC >0.5) were applied to the DEG lists. These lists were then used for downstream analysis.

Pathway analysis was performed using Metascape [https://metascape.org/] ^30^, a web-based platform that combines functional enrichment, gene annotation, interactome analysis and membership search from over 40 independent knowledgebases, within a single integrated portal. For pathway analysis, the genes that were up- or downregulated in *Nrp2*-KO macrophages, relative to expression in wild- type murine macrophages, were uploaded onto the Metascape platform and pathway enrichment was performed using the GO BP, KEGG Pathway and Reactome annotation memberships selected under the ‘enrichment’ tab. Pathways were considered enriched if they met the significance threshold of p<0.01, had a minimum overlap of 3 genes and had an enrichment score of >1.5. After running the enrichment analysis, Metascape generated a list of enriched pathways (ranked by enrichment score) as well as a table outlining the p-value, enrichment score and member genes for each enriched pathway. Further analysis and sorting of the genes within each enriched pathway was performed using Microsoft Excel. For more detailed visualisation of enriched pathways of interest, heatmaps of the member genes were generated using the online tool, Heatmapper [https://heatmapper.ca], and graphs of representative genes within the enriched pathway were generated in GraphPad Prism 7.0c, by plotting normalised DESeq2 gene counts.

### 2.15 Statistical analysis

Where possible, statistical analysis was performed on the data sets using GraphPad Prism 7.0c software. Values are expressed as the absolute mean ± standard error of the mean (SEM), as specified in the figure legends. Unless otherwise stated, analysis for statistical significance was performed blinded through unpaired, two-tailed student t-test or one-way analysis of variance (ANOVA) followed by Bonferroni post-hoc analysis, as specified in the figure legends, which have been reported as a p value followed by sample size. A statistically significant difference was defined as a result where p<0.05, unless otherwise stated.

## 3. Results

### 3.1 NRP2 expression during Macrophage Differentiation and Polarisation

Freshly harvested bone marrow cells were cultured in macrophage colony-stimulating factor (M-CSF)-enriched differentiation medium (30% L929-conditioned medium - LCM) for 7 days and macrophage differentiation was confirmed by assessment of cell morphology as well as mRNA and protein expression of the murine macrophage marker, F4/80 (Fig.S1). *Nrp2* mRNA was significantly upregulated from day 1 onwards in differentiating macrophages, reaching a maximum 13.6-fold increase relative to day 0 on day 5, and remaining significantly elevated at day 7 (7.9-fold increase in comparison to day 0) (Fig.1A). NRP2 protein could not be detected by western blot in cell lysates harvested directly after bone marrow extraction, or after 1 day of culture, indicating that early macrophage monocytic precursors and other cells in the bone marrow express very little NRP2 (Figure 1B) ^31^. After 3 days of culture, NRP2 protein was significantly upregulated before decreasing by 42% at day 5 and a further 24% by day 7 (Fig.1B,C). A similar pattern of NRP2 upregulation, followed by subsequent decline and plateauing, was reported previously in BMDMs differentiated using recombinant M-CSF ^23^, and may result from NRP2 expression in other non-macrophage cell types, such as osteocytes, skeletal muscle or adipocytes, present in cultures at day 3 but removed when the medium is changed at day 5 as part of the BMDMs differentiation protocol, thus decreasing total NRP2 protein expression.

**Figure 1.**
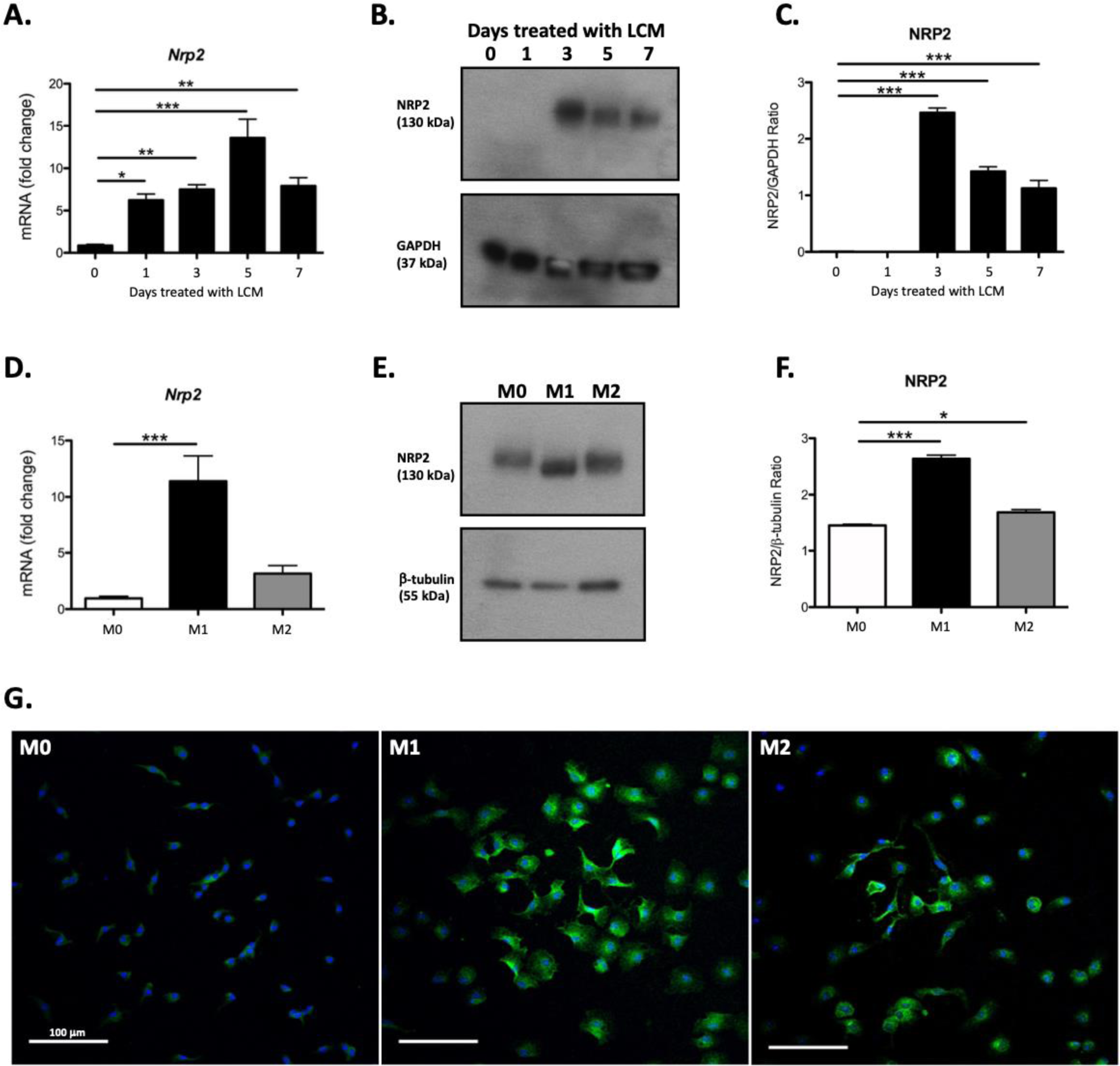
Expression of Neuropilin 2 throughout BMDM differentiation and in M1 and M2 polarised macrophages. **A.** Relative RT-qPCR analysis of *Nrp2* expression in BMDMs after 0, 1, 3, 5 and 7 days of treatment with 30% LCM. *Nrp2* expression was normalised against expression of β-actin and data is represented as mean fold-change relative to expression on day 0, with error bars showing S.E.M. (n=5). **B-C.** Representative Western Blot (**B**) and quantification graph (**C**) showing NRP2 (130 kDa) expression in BMDMs after 0, 1, 3, 5 and 7 days of treatment with 30% LCM. NRP2 expression was normalised against GAPDH (37 kDa) expression and data is represented as mean NRP2/GAPDH ratio, with error bars showing S.E.M (n=3). **F.** Relative RT-qPCR analysis of *Nrp2* expression in M0, M1 and M2 polarised macrophages after 24 hours of polarisation. Gene expression was normalised against *β-actin* expression and data is represented as mean fold-change relative to expression in M0 cells, with error bars showing S.E.M (n=5). **E-F.** Representative Western Blot (**E**) and quantification graph (**F**) showing NRP2 expression in M0 (serum only), M1 (LPS and IFNγ) and M2 (IL4) polarised macrophages after 48 hours of polarisation. NRP2 expression was normalised against β-tubulin (55 kDa) expression and data is represented as mean NRP2/β-tubulin ratio, with error bars showing S.E.M (n=3). **G.** Representative immunofluorescence images of M0, M1 and M2 polarised macrophages stained for NRP2 (green) and the nuclear stain, DAPI (blue) after 2 days of polarisation (n=3). * p<0.05, ** p<0.01, *** p<0.001 (one-way ANOVA with Bonferroni post-hoc analysis).

Changes in NRP2 expression were next determined during polarisation of differentiated BMDMs to either a pro-inflammatory M1-like state induced by treatment with LPS and Interferon (IFN)γ, or to an anti-inflammatory M2-like state, induced by treatment with Interleukin (IL)4. Treatment of BMDMs with LPS and IFNγ for 24 hours resulted in significant upregulation of mRNAs for well-characterised markers of M1 polarisation, including *Inos*, *Il1β*, *Tnfα* and *Il6*, with no significant increase in expression of the M2 polarisation markers *Arg1*, *Ym1* and *Cd206* (Fig.S2A). In line with gene expression, BMDMs treatment with LPS and IFNγ significantly increased iNOS expression (Fig.S2B), with no change in ARG1 protein levels (Fig.S2C). Further consistent with these findings, immunofluorescent staining demonstrated that, whereas iNOS could not be readily detected in untreated M0 macrophages, most cells stained positive for iNOS after 48h-treatment with LPS and IFNγ, whereas very few stained positive for ARG1 (Fig.S2D). Treatment with LPS and IFNγ resulted in significant *Nrp2* gene upregulation, with a 11.4-fold increase in *Nrp2* expression in M1 macrophages compared to M0 controls (Fig.1D). In line with gene expression, after 48 hours of treatment with LPS and IFNγ, NRP2 protein expression was significantly upregulated (Fig.1E,F). Interestingly, the apparent molecular weight of the NRP2 band detected in M1 macrophages lysates appeared marginally lower than that detected in M0 macrophages (Fig.1E), which may reflect distinct changes to the post-translational modifications of NRP2 following M1 polarisation. NRP2 post-translational modification by polysialylation has been documented in monocyte-derived mature dendritic cells, microglia, and peritoneal exudate macrophages ^31-33^. Werneburg et al. observed a shift in the apparent molecular weight of NRP2 in microglial cells after endosialidase treatment to remove polysialylation similar to that evident after LPS treatment in our study. They also reported that LPS stimulation of microglia caused complete depletion of polysialylation. It is therefore plausible that the LPS-induced shift of NRP2 to a lower molecular weight species found by us in macrophages could be due to an LPS-induced depletion of polysialylation similar to that reported in microglia. Immunofluorescent staining detected very little NRP2 expression in M0 macrophages, whereas cells stained positive for NRP2 after M1 polarisation, with a cytoplasmic and perinuclear distribution of the NRP2 protein (Fig.1G).

IL4- stimulated M2 polarisation resulted in the significant upregulation of the M2 markers, *Arg1* and *Cd206*, as well as a trend towards increased expression for *Ym1* (Fig.S2A). In line with gene expression, BMDMs treatment with IL4 significantly increased ARG1 expression (Fig.S2C), with no change in iNOS protein levels (Fig.S2B). Consistent with these findings, immunofluorescent staining demonstrated that IL4-treated cells stained positive for ARG1, whereas much less ARG1 staining was observed in untreated (M0) and M1 macrophages (Fig.S2D). IL4-treated cells also stained positive for iNOS, with immunostaining almost exclusively confined to the nucleus, as indicated by overlap of iNOS and DAPI staining (Fig.S2D).

Following 24h-treatment with IL4, *Nrp2* expression increased, but this did not reach statistical significance (Fig.1D). Following 48h-IL4 treatment, a modest yet significant increase in NRP2 expression was observed (Fig.1E,F). In line with western blotting data, NRP2 immunofluorescent staining also increased after M2 polarisation (Figure 1G).

In mammals, there are two distinct membrane-bound NRP2 isoforms, NRP2a and NRP2b, which share an identical extracellular domain but only share 11% sequence homology between their transmembrane and intracellular domains. Whilst NRP2a contains a PDZ binding motif at its C-terminus, as does NRP1, NRP2b does not, and thus has a slightly lower molecular weight than NRP2a. The two NRP2 isoforms have been proposed to have different roles, with NRP2b involved in the regulation of growth factor signalling and NRP2a thought to play a role in inflammation ^34^. Given the differences in overall NRP2 upregulation in M1 and M2 polarisation, and the shift in the molecular weight of NRP2 in M1 macrophages, we investigated whether *Nrp2a* and *Nrp2b* are differentially expressed following macrophage polarisation, by performing qPCR using primers specifically targeting non-homologous regions within the Nrp2a and Nrp2b C-terminus sequences. Following 24h-treatment with LPS and IFNγ, both *Nrp2a* and *Nrp2b* were significantly upregulated, however the magnitude of upregulation was strikingly different between the two Nrp2 isoforms (Fig.S3). *Nrp2a* expression was 11-fold greater in M1 macrophages relative to M0 macrophages (Fig.S3A), whereas *Nrp2b* expression increased by 47-fold (Fig.S3B). IL4-induced M2 polarisation caused a significant upregulation of *Nrp2a*, with a 5-fold increase in expression relative to M0 macrophages (Fig.S3A), whereas upregulation of *Nrp2b* in M2 macrophages was not statistically significant (Fig.S3B).

### 3.2 Nrp2 deletion regulates the expression of macrophage polarisation markers

To determine whether NRP2 is functionally important for macrophage polarisation, we investigated the impact of loss of NRP2 in BMDMs derived from a mouse line with myeloid-specific (LysM-Cre-restricted) deletion of *Nrp2* (*Nrp2-KO^Mac,EYFP^*) (Fig.S4A). *Nrp2-KO^Mac,EYFP^* mice were viable, produced litters with the expected Mendelian ratios, and exhibited no differences in size, weight or behaviour in comparison to wild-type (WT) littermates (data not shown), in line with previous reports^24^. *Nrp2* mRNA expression in BMDMs from *Nrp2-KO^Mac,EYFP^*mice (*LysM*-*Cre*^+/-^, *Nrp2^flx/flx^*, *EYFP^+/+^*) was reduced 9-fold in comparison to the expression in WT littermates (*LysM*-*Cre*^+/-^, *Nrp2^+/+^*, *EYFP^+/+^*) (Fig.S4C). Similarly, NRP2 protein expression was significantly reduced in BMDMs extracted from 10- to 12-week-old *Nrp2-KO^Mac,EYFP^* mice, with a 99.6% reduction in expression in comparison to WT littermates (Fig. S4D-E). With the added validation by EYFP expression in all observed cells (Fig.S4B), these results confirmed that Cre recombination had successfully occurred in macrophages, resulting in efficient knockout (KO) of NRP2 gene and protein.

In untreated M0 BMDMs from WT and *Nrp2-KO^Mac,EYFP^* mice, no significant difference in the expression of M1 and M2 polarisation markers were observed (Fig.2A-G). Following M1 polarisation with LPS and IFNγ, expression of the M1 markers, *Il1β*, *Tnfα* and *Il6* were all significantly lower in M1 polarised *Nrp2-KO^Mac,EYFP^* macrophages, compared to WT controls (Fig.2A-C). In contrast, expression of *Inos* was significantly increased in *Nrp2-KO^Mac,EYFP^*macrophages, compared with the WT control (Fig.2D). No significant difference in the expression of the M2 markers, *Arg1*, *Cd206* or *Ym1* were observed in M1 polarised *Nrp2-KO^Mac,EYFP^* macrophages in comparison to M1 polarised control cells (Fig.2E-G). Similarly, the expression of the M2 markers (*Arg1*, *Cd206* and *Ym1)* was not affected by the loss of NRP2 following M2 polarisation in *Nrp2-KO^Mac,EYFP^* macrophages compared with WT macrophages (Fig.2E-G).

**Figure 2.**
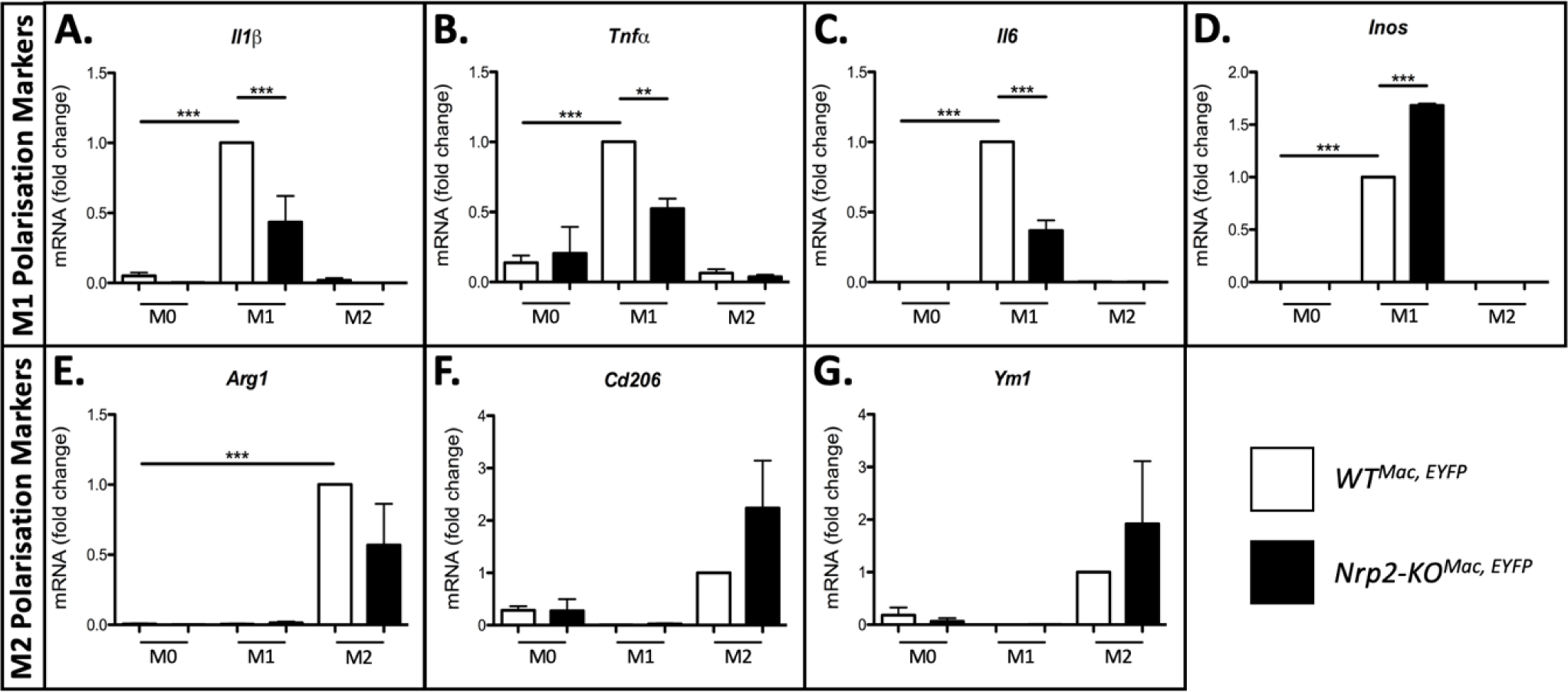
Impact of Neuropilin 2 knockout on macrophage polarisation. Relative RT-qPCR analysis of *Il1*β (**A**), *Tnf*α (**B**), *Il6* (**C**), *Inos* (**D**), *Arg1* (**E**), *Cd206* (**F**) and *Ym1* (**G**) expression in BMDMs extracted from *WT^Mac, EYFP^*(white bars) and *Nrp2-KO^Mac, EYFP^* (black bars) mice, following M0 (serum only), M1 (LPS and IFNγ) and M2 (IL4) polarisation. Gene expression was normalised against expression of β*-actin* expression and data represented as mean fold-changes relative to expression in *WT^Mac, EYFP^* M1 polarised macrophages (**A-D**), or *WT^Mac, EYFP^* M2 polarised macrophages (**E-G**), with error bars showing S.E.M (n=3). ** p<0.01, *** p<0.001 (one-way ANOVA with Bonferroni post-hoc analysis).

### 3.3 RNA-sequencing and pathway enrichment analysis in Nrp2-KO macrophages

The findings in Figure 2 indicated that loss of *Nrp2* had a predominantly inhibitory effect on pro-inflammatory gene expression markers but had no significant effect on markers of the anti-inflammatory M2 phenotype. To investigate further how NRP2 regulates gene expression in macrophages, whole transcriptome analysis was performed through RNA sequencing of BMDMs extracted from *Nrp2-KO^Mac,EYFP^* mice and WT controls. RNA-seq analysis revealed that *Nrp2* KO induces genome-wide changes in macrophage gene expression with 433 genes found to be downregulated, and 1274 genes upregulated, in BMDMs from *Nrp2-KO^Mac,EYFP^* mice relative to expression in WT controls (Fig.3A). The most significantly downregulated gene in *Nrp2*-KO macrophages was WD Repeat and FYVE domain containing 1 (*Wdfy1*) (L2FC=-0.826, Padj=7.39E-14) (Fig.3B), and the most significantly upregulated gene in these cells was microtubule associated protein tau (*Mapt*) (L2FC=7.161, Padj=4.21E-28) (Fig.3C). It is noteworthy that previous studies have implicated Wdfy1 in Toll-like receptor (TLR) 3/TLR4 immune responses and in the pathogenesis of inflammatory disease ^35-37^.

**Figure 3.**
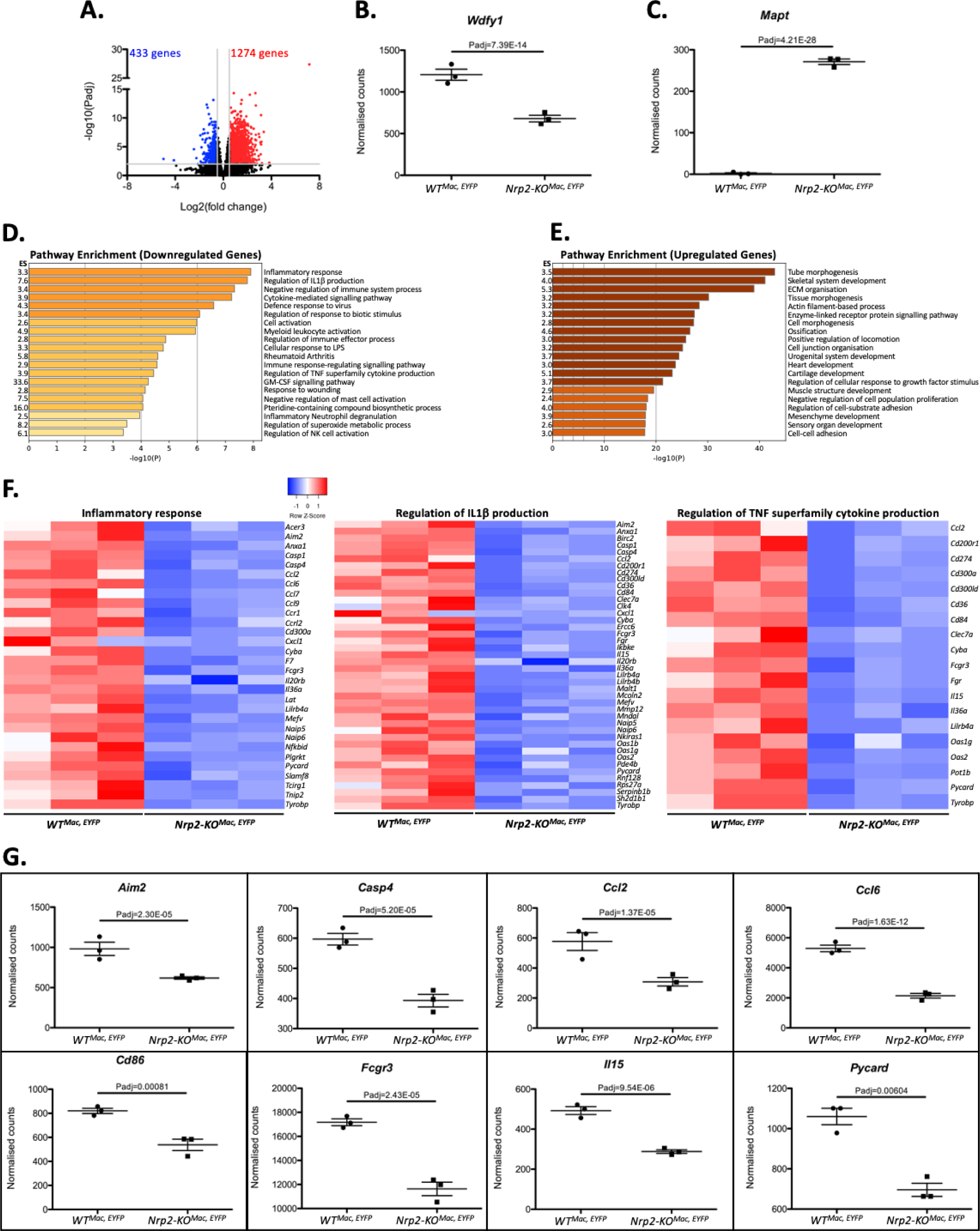
RNA-sequencing and pathway enrichment analysis in *Nrp2* knockout macrophages Downregulated and Upregulated genes in *Nrp2*-KO macrophages. **A.** Volcano plot showing the downregulated (blue) and upregulated (red) DEGs in *Nrp2*-KO (*Nrp2-KO^Mac, EYFP^*) macrophages, identified by RNA-seq. Grey lines represent the significance thresholds (-log10(Padj) >2 and -0.5> LFC >0.5), with non-DEGs in black. **B,C.** Graphs showing the normalised counts of the most significantly downregulated (**B**) and upregulated (**C**) genes from **A**. Graphs show the mean +/- S.E.M along with the Padj from DESeq analysis. **D,E.** Pathway Enrichment analysis of the downregulated (**D**) and upregulated (**E**) DEGs from **A** plotted by significance (-log10(P)), with enrichment scores (ES) included. Annotations refer to GO BP, KEGG Pathway and Reactome derived pathways. **F.** Heatmaps generated using the normalised counts of the downregulated DEGs contributing to the enrichment of the ‘inflammatory response’ (left), ‘regulation of IL1β production’ (centre) and ‘regulation of TNF superfamily cytokine production’ (right) pathways that were identified through RNA-seq analysis of *Nrp2*-KO macrophages (*Nrp2-KO^Mac, EYFP^*) and wild-type controls (*WT^Mac, EYFP^*). Colour scales indicate Z-score. **G.** Graphs showing the normalised counts of representative genes contained within the heatmaps shown in **F**. Graphs show the mean +/- S.E.M along with the Padj from DESeq2 analysis.

Enrichment analysis was performed on the up- and downregulated DEG lists, obtained from sequencing *Nrp2-KO^Mac,EYFP^* macrophages, using the GO BP, KEGG Pathway and Reactome annotation memberships on Metascape (Fig.3D,E). Enrichment analysis of the downregulated DEGs identified multiple NFκB-dependent pro-inflammatory pathways to be enriched including the ‘inflammatory response’, ‘regulation of IL1β production’, ‘cellular response to LPS’ and ‘regulation of TNF superfamily cytokine production’ pathways (Fig.3D). Enrichment analysis of the downregulated DEGs also identified a potential role for NRP2 in promoting migration since several of the genes contributing to NFκB-dependent pro-inflammatory pathway enrichment are also implicated in macrophage and monocyte migration. Heatmaps displaying the genes that contributed to the significant enrichment of the ‘inflammatory response’, ‘regulation of IL1β production’ and ‘regulation of TNF superfamily cytokine production’ pathways are shown in Fig.3F. The normalised DESeq2 counts for representative downregulated genes within the enriched pathways are shown in Fig.3G. A selection of inflammatory DEGs was further validated via RT-qPCR and confirmed a significant downregulation of *Pycard, Card9, Cyba, Ilb, Rab7 and Ikbk* in *Nrp2-KO^Mac,EYFP^* versus WT macrophages (Fig.S5).

Enrichment analysis of the upregulated DEGs identified growth factor receptor signalling pathways to be enriched, including several pathways related to extracellular matrix remodelling, TGFβ signalling and organogenesis. These enriched pathways include the ‘ECM organisation’, ‘heart development’ and ‘regulation of cell response to growth factor stimulus’ pathways (Fig.3E). Specific genes that were upregulated in *Nrp2-KO^Mac,EYFP^* versus WT macrophages were those involved in growth factor signalling (*Bcar1, Nedd4, Pdgfra, Pdgfrb, fgfr1, Egfr*), TGFβ family signalling (*Bmp5, Tgfb2*), extracellular matrix organisation (*Col11a1, Col3a1, Mmp2*), and Wnt signalling (*Wnt5a, Wnt5b, Yap1*). Interestingly, genes implicated in the Wnt signalling pathway were also found to contribute to the enrichment of multiple pathways, suggesting that NRP2 might play a role in the negative regulation of Wnt signalling in macrophages.

Overall, the findings from RNA-seq analysis of *Nrp2-KO^Mac,EYFP^* macrophages indicated that *Nrp2* ablation in macrophages impairs pro-inflammatory signalling pathways, consistent with our analysis of polarisation marker gene expression in these macrophages (Fig.2), showing that *Nrp2* KO impaired the upregulation of the pro-inflammatory cytokines, *Il1β*, *Tnfα* and *Il6,* following stimulation with LPS and IFNγ.

### 3.4 Myeloid-specific Nrp2-deletion reduces aortic plaque development in ApoE-KO mice fed a HFD

The effect of myeloid-specific *Nrp2* KO on atherosclerotic plaque development was determined by crossing *Nrp2-KO^Mac,EYFP^* to ApoE KO mice to generate *Nrp2-KO^Mac, Apoe-/-,EYFP^* mice, and examining the effect of feeding these mice a high fat diet (HFD) for 16 weeks as compared to *WT^Mac,Apoe-/-,EYFP^*controls. Prior to commencing HFD, there was no significant difference in the weights of *Nrp2-KO^Mac, Apoe-/-, EYFP^* and control *WT^Apoe-/-, EYFP^* mice (Fig.S6A), and no significant difference in weights was noted after 16 weeks of HFD (Fig.S6A). Similarly, there was no change in the weights of the liver, spleen or kidneys (Fig.S6B-D) between *Nrp2-KO^Mac, Apoe-/-,EYFP^* and control mice.

Oil-red-O staining of aortas showed that *Nrp2-KO^Mac, Apoe-/-,EYFP^* mice developed significantly less total aortic plaque compared to *WT^Apoe-/-, EYFP^* controls (Fig.4A), with 17.4% of the aortas staining positive for Oil-red-O in *Nrp2-KO^Mac, Apoe-/-,EYFP^* mice, compared to 24.9% in control mice (Fig.4B). Plaque coverage within both the aortic arch and the thoracic aorta was significantly lower in *Nrp2-KO^Mac, Apoe-/-,EYFP^* mice (Fig.4B). A trend towards reduced plaque coverage within the abdominal aorta was also observed in *Nrp2-KO^Mac, Apoe-/-,EYFP^* mice, but was not statistically significant (Fig.4B). Next, we assessed aortic root sections for plaque characteristics that are important markers of plaque stability in human atherosclerosis ^38-41^, including cap thickness, necrotic core area, lipid content, and macrophage and smooth muscle cell (SMC) recruitment (Fig.5 and Fig.S7). Lipid content in aortic roots of *Nrp2-KO^Mac, Apoe-/-,EYFP^* mice was decreased in comparison to controls, with Oil-red-O staining covering 11.5% of total plaque in *Nrp2-KO^Mac, Apoe-/-,EYFP^*mice, compared to 19.4% in *WT^Apoe-/-, EYFP^* mice (Fig.5A-C). A trend towards reduced total aortic root plaque area was also observed *Nrp2-KO^Mac, Apoe-/-,EYFP^* mice compared to control mice (Figure 5A’-C’), though this was not statistically significant. Plaque cap was significantly thicker (52.8% increase) in *Nrp2-KO^Mac, Apoe-/-, EYFP^* mice compared to control mice (average cap thickness of 22.14 μm in *Nrp2-KO^Mac, Apoe-/-,EYFP^* mice versus 14.49 μm in controls) (Fig.5A’’-C’’). In contrast, the necrotic core was significantly smaller in aortic root plaque in *Nrp2-KO^Mac, Apoe-/-,EYFP^* mice, with the necrotic core accounting for 23.9% of total plaque area in *Nrp2-KO^Mac, Apoe-/-,EYFP^* mice, compared to 33.8% in *WT^Apoe-/-,EYFP^*control mice (Fig.5A’’’-C’’’).

**Figure 4.**
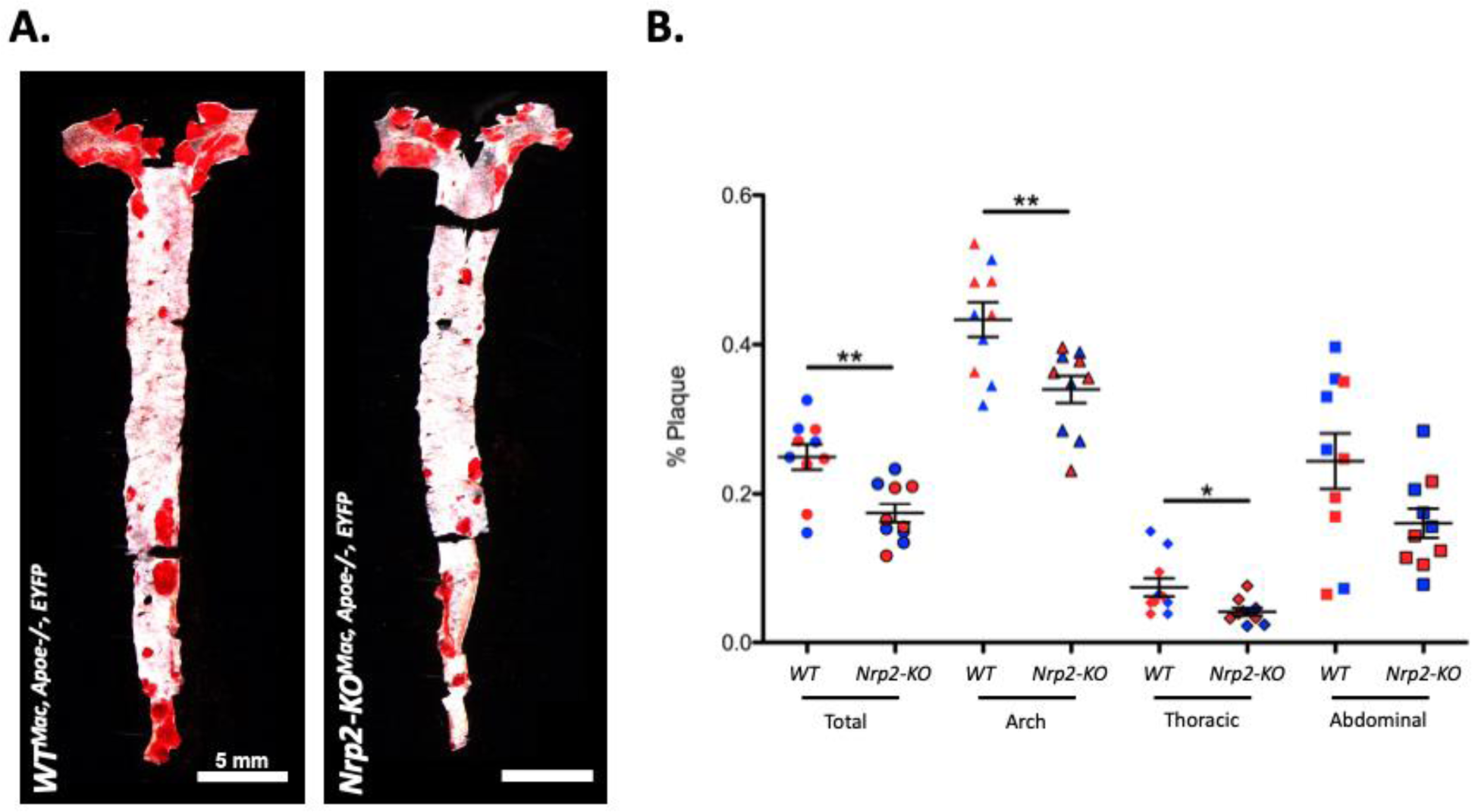
Impact of myeloid-specific Neuropilin 2 knockout on high-fat diet-induced aortic plaque development. Representative Oil-red-O-stained whole aortas (**A.**) and percentage plaque quantification graph (**B.**) from *WT^Mac, Apoe-/-, EYFP^* (*WT*) and *Nrp2-KO^Mac, Apoe-/-, EYFP^* (*Nrp2-KO*) mice after 16 weeks of HFD feeding (5 mm scale bar). **B.** Graph shows mean percentage plaque within the total aorta and within segmented regions of the aorta (arch, thoracic, abdominal) +/- S.E.M (n=10), with male (blue) and female (red) mice indicated. The posterior intercostal arteries and the mesenteric arteries were used to define the thoracic and abdominal aorta, respectively. * p<0.05, ** p<0.01 (unpaired two-tailed t-test).

**Figure 5:**
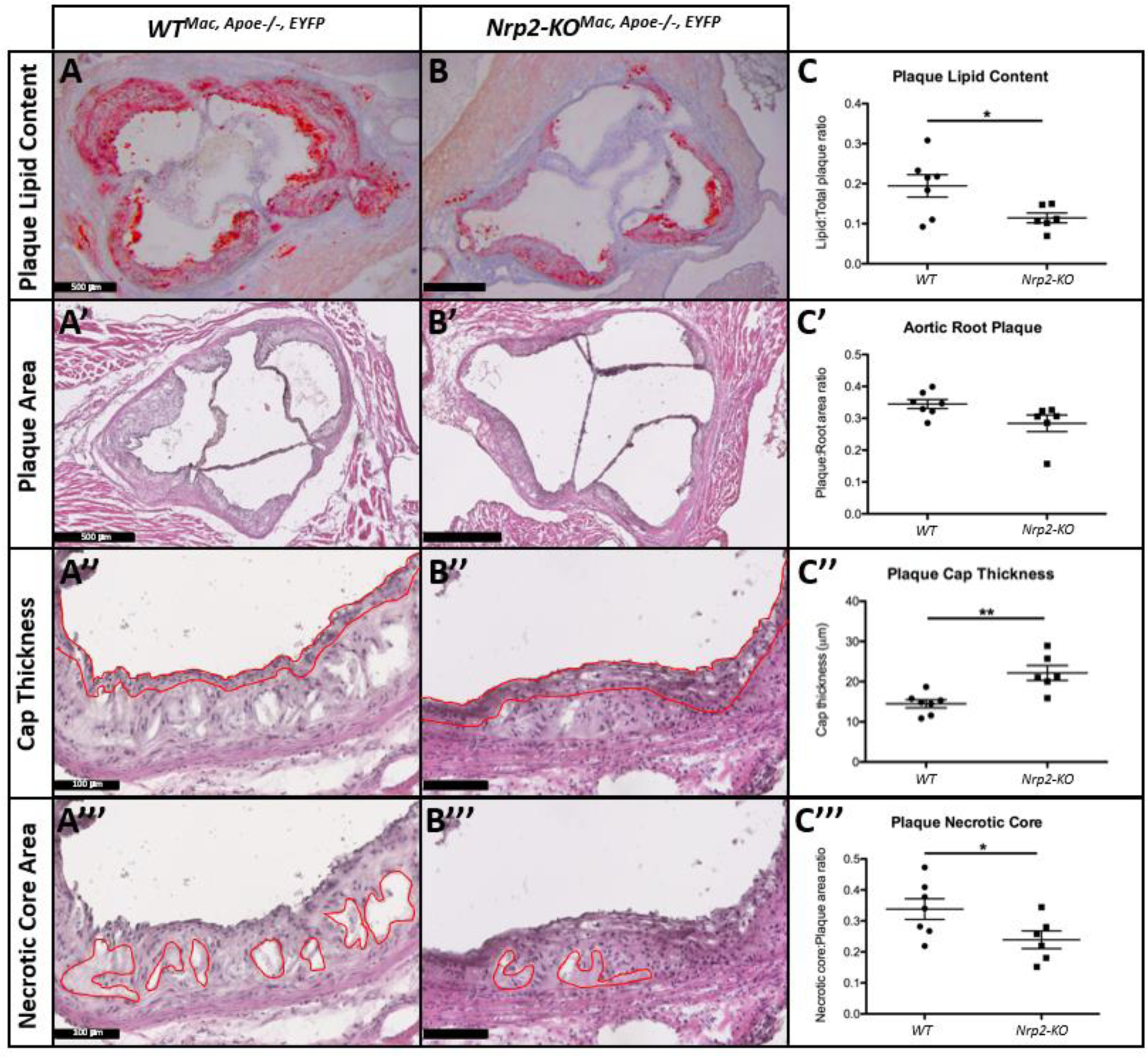
Impact of myeloid-specific Neuropilin 2 knockout on high-fat diet-induced aortic plaque size, cap thickness, necrotic core area and lipid content. **A-B.** Representative images (500 μm scale bar) of plaque lipid content in Oil-red-O-stained aortic root sections from *WT^Mac, Apoe-/-, EYFP^* (left) and *Nrp2-KO^Mac, Apoe-/-, EYFP^* (right) mice after 16 weeks of HFD feeding. **A’-B’’’.** Representative images of plaque area (**A’-B’**, 500 μm scale bar), cap thickness (**A’’-B’’**, 100 μm scale bar, cap defined by red area) and necrotic core area (**A’’’-B’’’**, 100 μm scale bar, necrotic core defined by red area) in H&E-stained aortic root sections from *WT^Mac, Apoe-/-, EYFP^* (left) and *Nrp2-KO^Mac, Apoe-/-, EYFP^* (right) after 16 weeks of HFD feeding. Graphs (**C-C’’’**) show the quantified mean relative lipid area (**C**), relative plaque area (**C’**), cap thickness (μm) (**C’’**) and relative necrotic core area (**C’’’**) +/- S.E.M (n>6) in *WT^Mac, Apoe-/-, EYFP^* (*WT*, circles) and *Nrp2-KO^Mac, Apoe-/-, EYFP^* (*Nrp2-KO*, squares) mice. Each datapoint represents the average quantified area or cap thickness across 2-3 aortic root sections per mouse. * p<0.05, ** p<0.01 (unpaired two-tailed t-test).

Plaque macrophage and SMC content were examined using immunofluorescent YFP expression to identify macrophages and an anti-αSMA antibody to detect SMCs. YFP expression was observed throughout the aortic root plaque (Fig.S7A,B), whereas αSMA expression was predominantly confined to the tunica media and the plaque cap (Fig.S7A’,B’). The overall intensity of YFP staining was similar in *Nrp2-KO^Mac, Apoe-/-,EYFP^* and *WT^Apoe-/-,EYFP^*mice. However, a substantial region of αSMA expression was observed at the luminal surface of aortic root plaques in *Nrp2-KO^Mac, Apoe-/-, EYFP^* mice (Fig.S7B’), which was not observed in control mice (Fig.S7A’). Furthermore, in contrast to control mice, considerable overlap in αSMA and YFP expression was observed within the plaque core in aortic root plaques from *Nrp2-KO^Mac, Apoe-/-,EYFP^* mice (compare Fig.S7A’’ and B’’). In *Nrp2- KO^Mac, Apoe-/-,EYFP^* plaques, αSMA expression was detected deep within the plaque core and could indicate a phenotypic switch of the macrophages towards an SMC-like profile.

### 3.5 Nrp2 is required for macrophage migration

Transcriptomic analysis and validation of DEGs via qPCR demonstrated that genes encoding chemokines, chemokine receptors and other molecules important for macrophage migration and chemotaxis pathways, such *as Ccl2, Ccl6, Ccl7, Cxcr2, Anxa1, Lgals3* and *Ccr1*, were significantly downregulated in *Nrp2-KO^Mac,EYFP^* macrophages (Fig.6A,B), therefore we next investigated whether the loss of *Nrp2* was functionally significant using a transwell chemotactic assay. Since *Ccl2*, encoding Monocyte Chemoattractant Protein-1 (MCP-1), was significantly downregulated in *Nrp2- KO^Mac,EYFP^* macrophages, and is a key chemoattractant in monocyte recruitment in atherogenesis, we investigated the effect of *Nrp2* KO on the macrophage migratory response to MCP-1. As shown in Fig. 6C, MCP-1-stimulated macrophage migration was significantly reduced in *Nrp2-KO^Mac,EYFP^*macrophages, with a 44.3% reduction relative to WT controls. Similar results were obtained when MCP-1 stimulation of *Nrp2-KO^Mac, Apoe-/-,EYFP^* macrophage migration was examined (data not shown).

**Figure 6:**
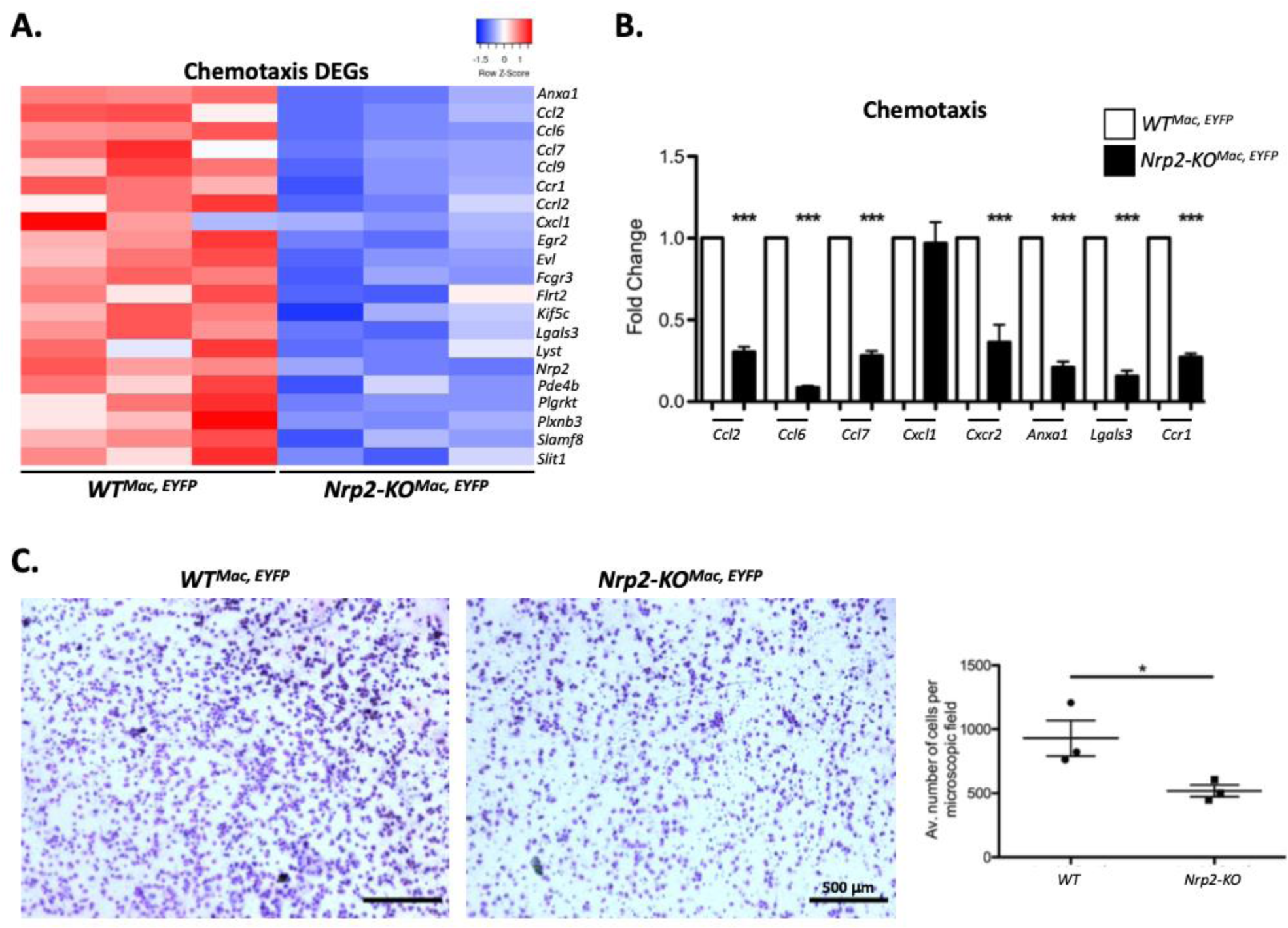
Impact of Neuropilin 2 knockout on macrophage migration towards MCP-1 and validation of chemotactic differentially expressed genes in Neuropilin 2 knockout macrophages. **A.** Heatmap generated using the normalised counts of the downregulated DEGs contributing to the enrichment of the ‘Chemotaxis’ pathway that were identified through RNA-seq analysis of *Nrp2-KO^Mac, EYFP^* macrophages and *WT^Mac, EYFP^* controls. Colour scales indicate Z-score. **B.** Relative RT-qPCR analysis of ‘Chemotaxis’ pathway gene expression in BMDMs extracted from *WT^Mac, EYFP^* (white bars) and *Nrp2-KO^Mac, EYFP^* (black bars) mice. Gene expression was normalised against expression of *β-actin* expression and data for each gene are represented as mean fold-change relative to expression in *WT^Mac, EYFP^* controls, with error bars showing S.E.M (n=3). *** p<0.001 (unpaired two-tailed t-test). **C.** Representative images of transwell migration assay membranes and quantification graph from *WT^Mac, EYFP^* (*WT*, left) and *Nrp2-KO^Mac, EYFP^*(*Nrp2-KO*, right) macrophages after 20 hours of migration towards MCP-1. Data are represented as the average number of cells per microscopic field, across 5 microscopic fields per transwell membrane, with graph showing mean +/- S.E.M (n=3). * p<0.05 (unpaired two-tailed t-test).

## 4. Discussion

In accord with previous studies, we found that NRP2 expression is very low or undetectable in bone marrow-derived mouse monocytes, and its expression increases during monocyte to macrophage differentiation (27). However, previous studies have been unclear as to NRP2 expression during macrophage polarisation. Here we demonstrate that NRP2 is markedly upregulated following pro-inflammatory activation of BMDMs by LPS and IFNγ, whereas anti-inflammatory polarisation in response to IL4 had little significant effect on NRP2 expression. Similar to our results in BMDMs, NRP2 expression increased during LPS-induced maturation of dendritic cells ^42^. In contrast, Ji et al. reported that LPS reduced NRP2 expression in human monocytes ^25^, but the methodology in this paper diverged significantly from ours as they only examined the effect of LPS alone rather than the combination of LPS and IFN-γ, and monocytes were treated following only 1 day of culture with M-CSF, in contrast to the 7 days treatment of mouse bone marrow cells used here and elsewhere ^26^. A study of NRPs in tissue-specific alveolar macrophages, found NRP1 and NRP2 expression in all human alveolar macrophages examined, but did not find conclusive evidence for a preferential association of either NRP with any specific macrophage subtype based on immunohistochemical analysis ^43^. Ours is therefore the first study to find compelling evidence for selective NRP2 upregulation in pro-inflammatory macrophage polarisation. We further showed that myeloid-restricted NRP2 deletion inhibited expression of several cytokine markers of inflammatory macrophage polarisation and caused a transcriptome-wide suppression of several key pathways in inflammation, including NFκB activation and monocyte/macrophage chemoattraction, fundamental mechanisms mediating inflammatory processes important for atherogenesis. These findings together provide strong evidence that NRP2 is required for inflammatory activation of macrophages.

An important role for NRP2 in facilitating NFκB signalling in macrophages is consistent with reduced expression of *Il1β*, *Tnfα* and *Il6,* all established NFκB pathway target genes, and by enrichment of NFκB-dependent pathways in the downregulated DEGs in *Nrp2-KO^Mac,EYFP^* macrophages. The most highly enriched pathways within the downregulated DEGs identified in *Nrp2*-KO macrophages were ‘inflammatory response’, and ‘regulation of IL1β production’, while ‘cytokine-mediated signalling pathway’, ‘cellular response to LPS’ and ‘regulation of TNF superfamily cytokine production’ were also enriched pathways in this gene set, all pathways in which NFκB is centrally involved. Gene-specific qPCR analysis showed that among the ‘inflammatory response’ pathway genes, *Iκbκb*, encoding inhibitor of κB kinase β (Iκκβ), was strongly downregulated in *Nrp2*-depleted macrophages. Iκκβ is essential for NFκB activation through phosphorylation of IκB causing its subsequent degradation.

In contrast to NRP1, evidence for a contribution of NRP2 to cell migration or chemotaxis is more sparse, and little is currently known about the role of NRP2 in macrophage migration ^44, 45^. However, NRP2 has been implicated in the migration of dendritic cells, through NRP2 polysialylation and binding of the polysialic acid moiety to the CCL21 chemokine and enhancement of its binding to CCR7 ^42^. The argument that NRP2 mediates macrophage migration is supported in the present study by the reduced expression in *Nrp2-KO^Mac,EYFP^* macrophages of several chemokine and chemokine receptors with essential roles in monocyte/macrophage migration and plaque recruitment, including *Ccl2*/MCP-1. Migration of *Nrp2-KO^Mac,EYFP^* macrophages towards MCP-1 was also impaired compared with wild-type cells indicating that deletion of *Nrp2* also inhibited the mechanism distal to MCP-1 cell binding. These findings also lend further weight to the conclusion that NRP2 is important for NFκB signalling, since NFκB induces macrophage chemotaxis in part through increased expression of MCP-1 and downstream mechanisms.

Our findings are in agreement with previous studies in tumour-associated macrophages which show both that elevated expression of NRP2 is associated with high expression of transcripts for inflammatory cytokines and chemokines, including TNFα, IL6, and CCL2, and that genetic depletion of the NRP2b isoform using stable expression of shRNA in the mouse macrophage RAW264.7 cell line markedly reduced IL6 expression ^46^. We found that NRP2b was more strikingly upregulated in response to LPS & IFNγ-induced inflammatory macrophage polarisation compared with NRP2a, and was not significantly increased in response to IL-4, suggesting that the NRP2b isoform is primarily responsible for the regulation of inflammatory signalling pathways revealed by RNAseq analysis.

It is of interest that the most strongly down-regulated gene in *Nrp2-KO^Mac,EYFP^* macrophages identified in our RNAseq analysis was the WD repeat and FYVE-domain-containing 1 protein (Wdfy1). Wdfy1 is a critical adapter molecule in TLR3/TLR4 signalling ^35^, a pathway which is important for diverse immune responses, and for the pathogenesis of inflammatory diseases ^36, 37, 47^. TLRs, especially TLR4, are also increasingly implicated in the pathogenesis of atherosclerosis ^48^. Though it was not within the scope of the present study to examine further the role of Wdfy1 in NRP2-mediated TLR3/TLR4 signalling in macrophages, our findings are consistent with an important role for NRP2 in TLR signalling relevant for pro-atherogenic inflammatory signalling. Another striking feature of gene expression in *Nrp2-KO^Mac,EYFP^* macrophages was the downregulation of multiple components of the NLRP3 inflammasome (*Pcard, Ilb*, and *Aim2*), a key driver of atherogenesis. Other down-regulated genes identified in *Nrp2-KO^Mac,EYFP^*macrophages by RNAseq and targeted qPCR included those essential for macrophage efferocytosis (*Rab7*), and pro-inflammatory cell death or pyroptosis (*Casp4*) ^49, 50^.

In marked contrast to the enrichment of inflammatory signalling pathways in genes downregulated by *Nrp2* deletion in macrophages, upregulated DEGs were enriched in pathways related to extracellular matrix remodelling, the cell response to growth factors, and tissue morphogenesis including genes involved in TGFβ and other growth factor signalling pathways, the Wnt pathway, and collagens. Several of the specific genes in these pathways that were upregulated in *Nrp2-KO^Mac,EYFP^* macrophages have been shown to promote M2 macrophage polarisation, including *Tgfb, Wnt5a, and Col3a1* ^51-58^. It is possible that the upregulation of genes within these pathways in *Nrp2*-KO macrophages is secondary to downregulation of M1-like inflammatory pathways rather than a direct result of NRP2 negatively regulating these pathways, though further work is necessary to elucidate the interactions between these diverse pathways. However, taken together, the results of RNAseq analysis suggest that *Nrp2* ablation in macrophages reprogrammes the inflammatory/anti-inflammatory axis within BMDMs, down-regulating pro-inflammatory pathways, and upregulating anti-inflammatory pathways associated with tissue growth and repair.

In contrast to the findings presented here for NRP2, there is compelling evidence that NRP1 suppresses pro-inflammatory macrophage functions ^18^. Thus, myeloid-restricted deletion of *Nrp1* inhibits the immunosuppressive activity of mouse TAMs thereby decreasing tumour growth and metastasis, enhances inflammatory cytokine production via TLR4 and NF-κB, and increased insulin resistance via NLRP3 inflammasome priming and activation ^19-22^. The present study indicates that NRP1 and NRP2 play divergent roles in macrophage function, broadly consistent with the hypothesis that NRP1 inhibits, whereas NRP2 promotes, activation of inflammatory pathways in macrophages. This conclusion is supported by our unpublished findings showing that in marked contrast with NRP2, NRP1 mRNA and protein were markedly downregulated in BMDMs during M1 polarisation and enhanced during M2 polarisation, and that genetic deletion of *Nrp1* in macrophages increased expression of the M1 polarisation markers, *Tnfα*, *Inos*, *Il1β* and *Il6*. The present study indicates that functions of NRP2 in inflammatory macrophages are important both for atherosclerotic plaque generation and in the formation of key determinants of plaque instability. This conclusion is supported by our data showing that myeloid-specific ablation of *Nrp2* significantly impaired total, aortic arch and thoracic aorta plaque formation in *Apo E^-/-^* mice. Furthermore, the significant increase in plaque cap thickness and concomitant decrease in both plaque necrotic core area and lipid content in *Nrp2-KO^Mac, Apoe-/-,EYFP^* mice indicates that loss of macrophage-derived NRP2 is likely to enhance plaque stability, a possibility also consistent with the observation that SMC were more prevalent in *Nrp2-KO^Mac, Apoe-/-,EYFP^* plaques. The mechanisms underpinning the inhibitory effect of myeloid-specific *Nrp2* ablation on atherosclerosis are likely mediated by changes in gene expression revealed by transcriptomic and targeted qPCR analysis of *Nrp2-KO^Mac,EYFP^* macrophages. Indeed, several of the downregulated genes in *Nrp2-KO^Mac,EYFP^* macrophages have known pro-atherogenic roles in atherosclerosis, including *Pycard*, also known as ASC (Apoptosis-associated speck-like protein containing a CARD), *Il1β*, and *Aim2*, all components of the NLRP3 inflammasome required for atherogenesis in *Ldlr*-deficient mice ^59, 60^. Also downregulated in *Nrp2-KO^Mac,EYFP^* macrophages was *Fcgamma3* (CD16), loss of which reduces lesion size in *Ldlr*-deficient mice ^61^ due to reduced pro-inflammatory gene expression, nuclear factor-κB activity, and M1 macrophage polarisation ^62^. Conversely, several of the enriched pathways that were upregulated in *Nrp2-KO^Mac,EYFP^*macrophages, such as matrix reorganisation, growth factor and Wnt signalling are linked to tissue growth and repair, predicted to have a generally anti-inflammatory effect and contribute to increased plaque stability. Overall, the transcriptomic signature of *Nrp2*-depleted macrophages is anti-atherogenic, consistent with a role of NRP2 in the pathogenesis of atherosclerosis. In line with this conclusion is a recent study reporting that tail vein injection of Adeno-associated virus (AAV) encoding NRP2-targeted RNAi into ApoE KO mice reduced lesion formation caused by high cholesterol diet feeding ^63^. An atherogenic role of NRP2 is also consonant with previous work showing that NRP2 knockdown using targeted shRNA significantly reduced neointimal hyperplasia following arterial injury, a process important for development of atherosclerotic plaques ^44^. These findings suggest that NRP2 may be a possible future target for inhibition of atherosclerotic plaque formation. Further elucidation of the mechanisms underpinning NRP2-mediated pro-inflammatory macrophage activation is also likely to improve understanding of the central role of macrophages in atherosclerotic disease.

## Acknowledgments

We would like to thank the UCL BSU Staff for help with husbandry and UCL Genomics for RNA sequencing services and pipeline analysis. The *R26R-EYFP*^+/+^, *ApoE*^−/−^ mouse line was a generous gift from the lab of Prof. Gary Owens’ Laboratory (Virginia, USA).

## Supplementary Tables

**Supplementary Table I:**
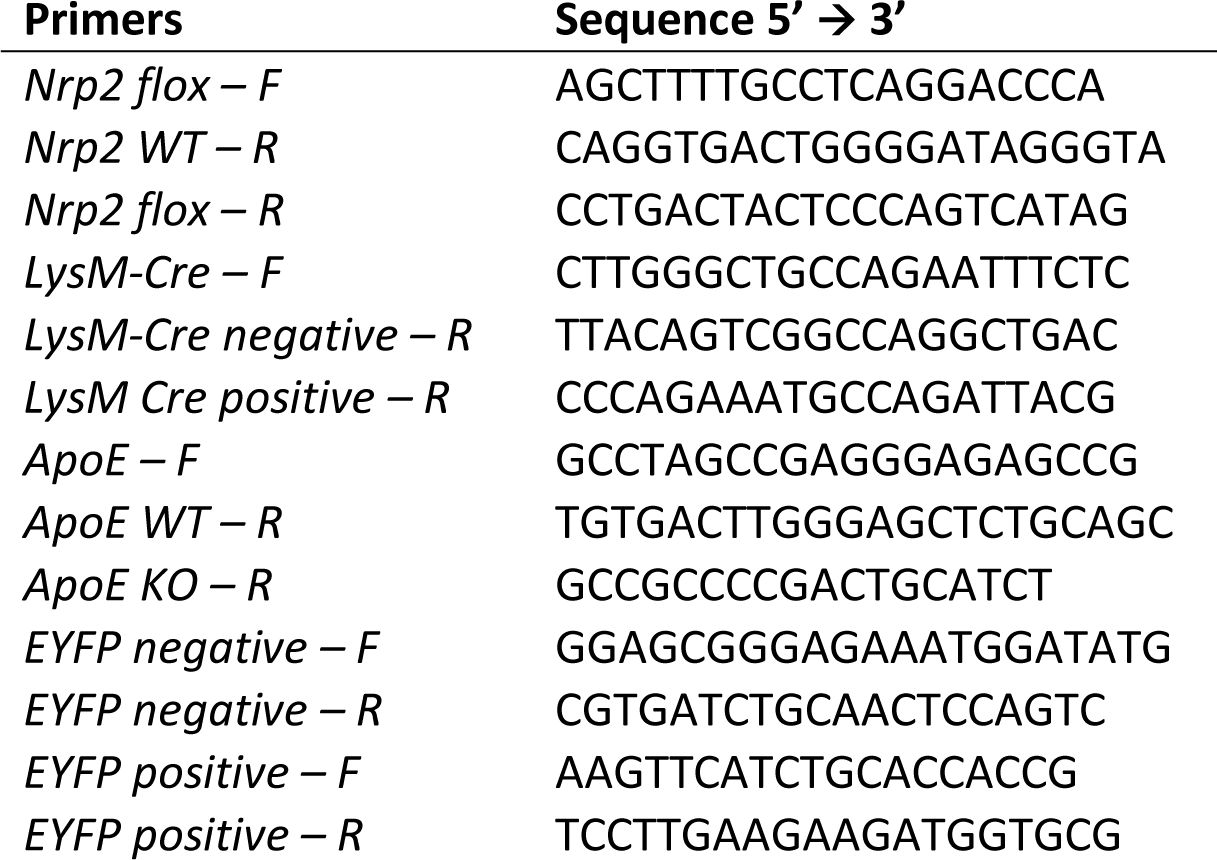
Genotyping primers.

**Supplementary Table II:**
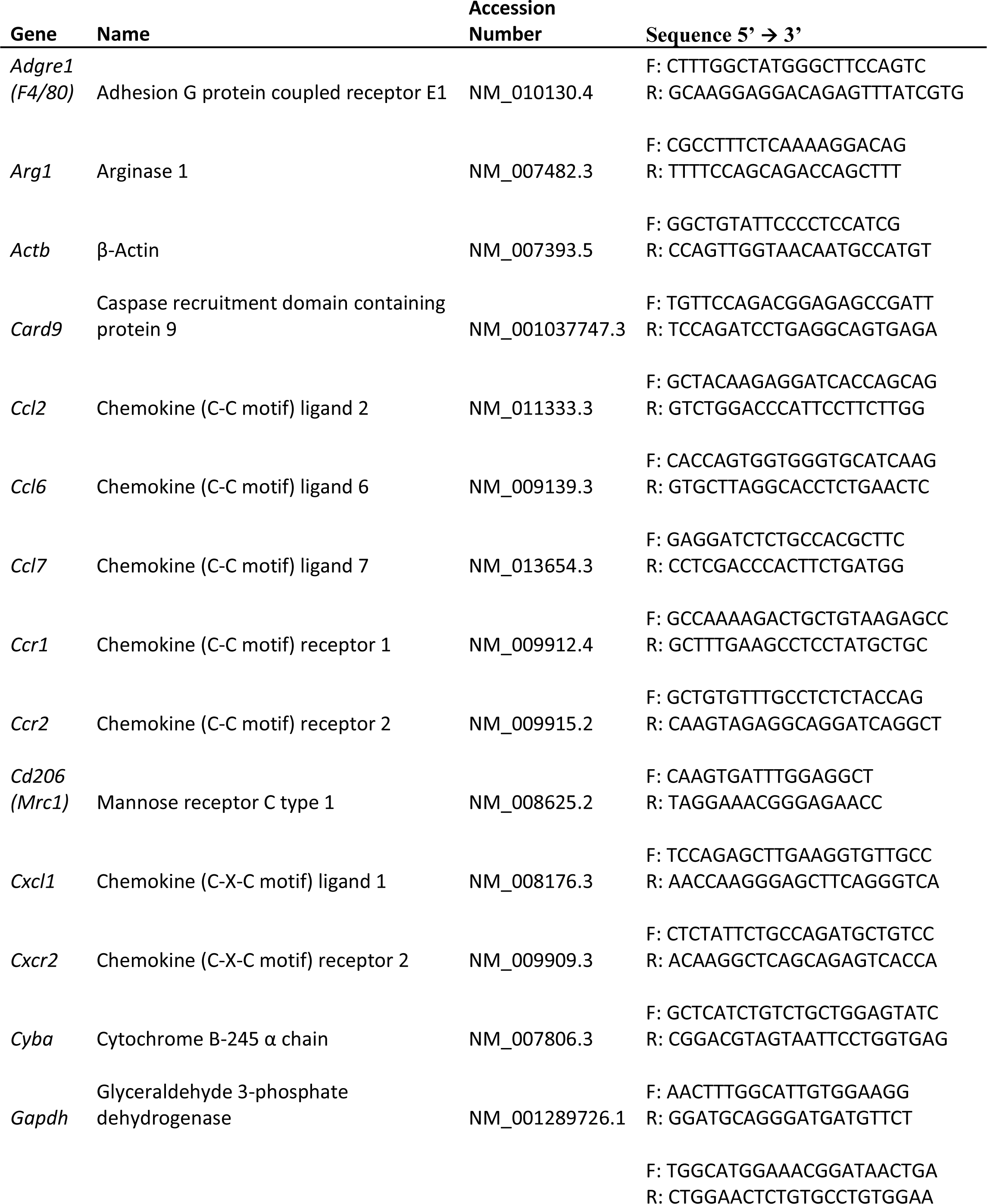
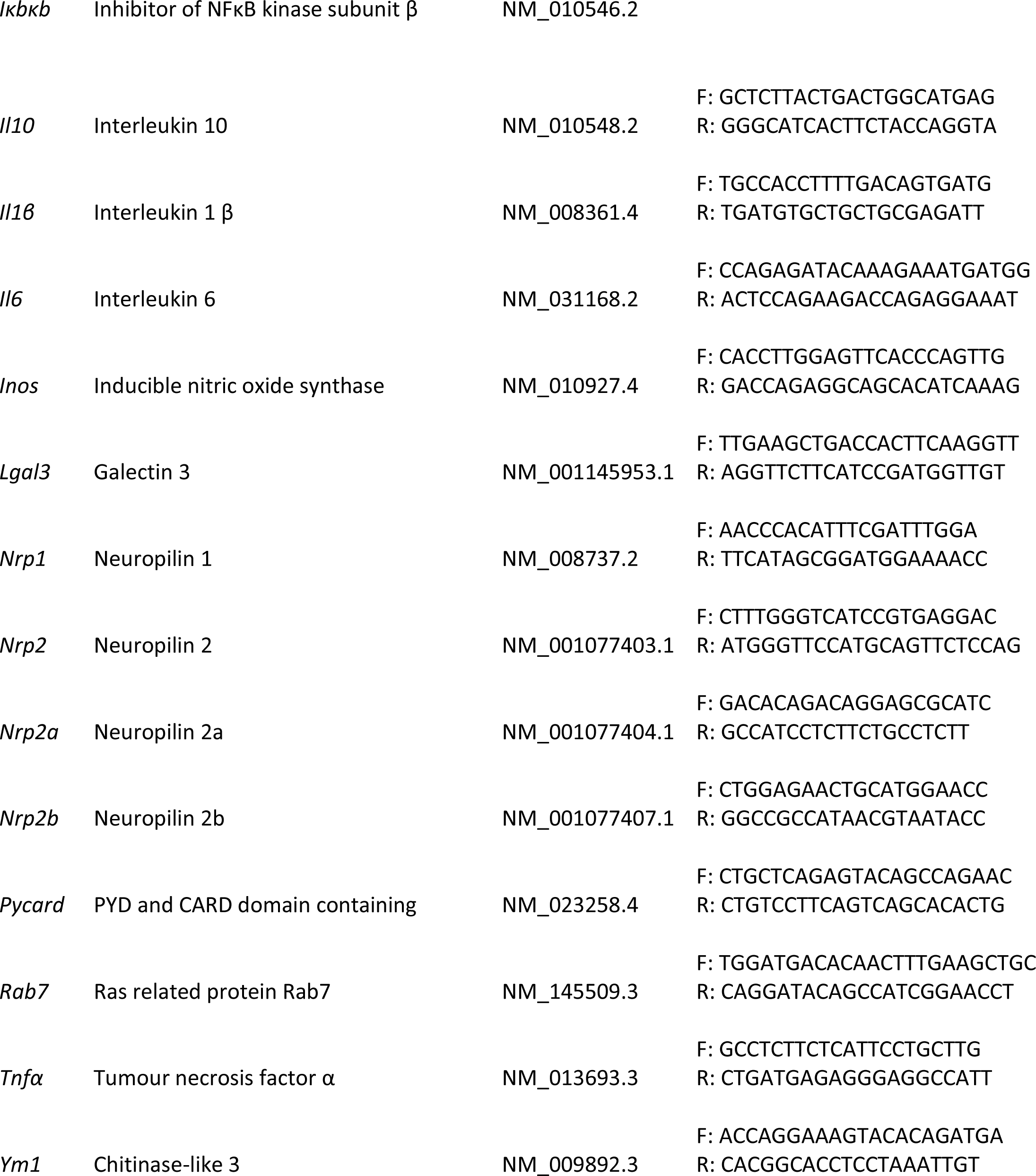
RT-qPCR primers.

**Supplementary Table III:** Antibodies.

| **Antibody Name** | **Antigen Labelled** | **Manufacturer** | **Catalogue number** | **Dilution** | **Application** |
| --- | --- | --- | --- | --- | --- |
| GAPDH (6C5) | Glyceraldehyde 3-phosphate dehydrogenase | Santa Cruz Biotechnologies | SC32233 | 1:500 | Western blot |
| β-tubulin (G-8) | β-tubulin | Santa Cruz Biotechnologies | SC55529 | 1:500 | Western blot |
| HSP90 (C45G5) | Heat shock protein 90 | Cell Signalling | #4877T | 1:500 | Western blot |
| Neuropilin-1 (EPR3113) | NRP1 C-terminus | Abcam | ab81321 | 1:500 | Western blot |
| Neuropilin-2 (D39A5) XP | NRP2 extracellular domain | Cell Signalling | #33665 | 1:500 | Western blot |
| iNOS/NOS type II | Inducible nitric oxide synthase | BD Biosciences | 610332 | 1:500 | Western blot |
| Liver Arginase (ARG1) | Liver arginase (arginase 1) | Abcam | ab60176 | 1:500 | Western blot |
| HRP-conjugated horse antimouse IgG | Mouse IgG | Cell Signalling | #7076 | 1:10000 | Western blot |
| HRP-conjugated goat anti-rabbit IgG | Rabbit IgG | Cell Signalling | 7074P2 | 1:10000 | Western blot |
| HRP-conjugated donkey antigoat IgG | Goat IgG | Cell Signalling | SC2020 | 1:10000 | Western blot |
| Neuropilin-1 (EPR3113) | NRP1 C-terminus | Abcam | ab81321 | 1:100 | Fluorescent ICC |
| Neuropilin-2 (D39A5) XP | NRP2 extracellular domain | Cell Signalling | #33665 | 1:100 | Fluorescent ICC |
| iNOS/NOS type II | Inducible nitric oxide synthase | BD Biosciences | 610332 | 1:100 | Fluorescent ICC |
| Liver Arginase (ARG1) | Liver arginase (arginase 1) | Abcam | ab60176 | 1:100 | Fluorescent ICC |
| NF-κB p65 (RelA) | Nuclear factor kappa B - p65 (RelA) | Cell Signalling | 8242 | 1:100 | Fluorescent ICC |
| Rat anti-mouse F4/80: Alexa Fluor® 488 | F4/80 antigen | Bio-Rad | 108551 | 1:100 | Fluorescent ICC |
| Alexa Fluor® 488 donkey anti-goat IgG (H+L) | Goat IgG | ThermoFisher Scientific | A11055 | 1:500 | Fluorescent ICC |
| Alexa Fluor® 488 goat anti-rabbit IgG (H+L) | Rabbit IgG | ThermoFisher Scientific | A11034 | 1:500 | Fluorescent ICC |
| Alexa Fluor® 488 donkey anti-mouse IgG (H+L) | Mouse IgG | ThermoFisher Scientific | A32723 | 1:500 | Fluorescent ICC |
| Anti-GFP protein Mouse IgG1κ | GFP and YFP | Roche | 11814460001 | 1:100 | Fluorescent IHC |
| αSMA [1A4] (Alexa Fluor® 594) | Alpha smooth muscle actin | Abcam | Ab202368 | 1:100 | Fluorescent IHC |
| Alexa Fluor® 488 rabbit anti-mouse IgG (H+L) | Mouse IgG | Molecular Probes | A21204 | 1:500 | Fluorescent IHC |
| Alexa Fluor® 594 mouse IgG2a, κ monoclonal [10D7A7B2] – isotype control | Mouse IgG | Mouse IgG | Ab178001 | 1:500 | Fluorescent IHC |

## Supplementary figures

**Figure S1.**
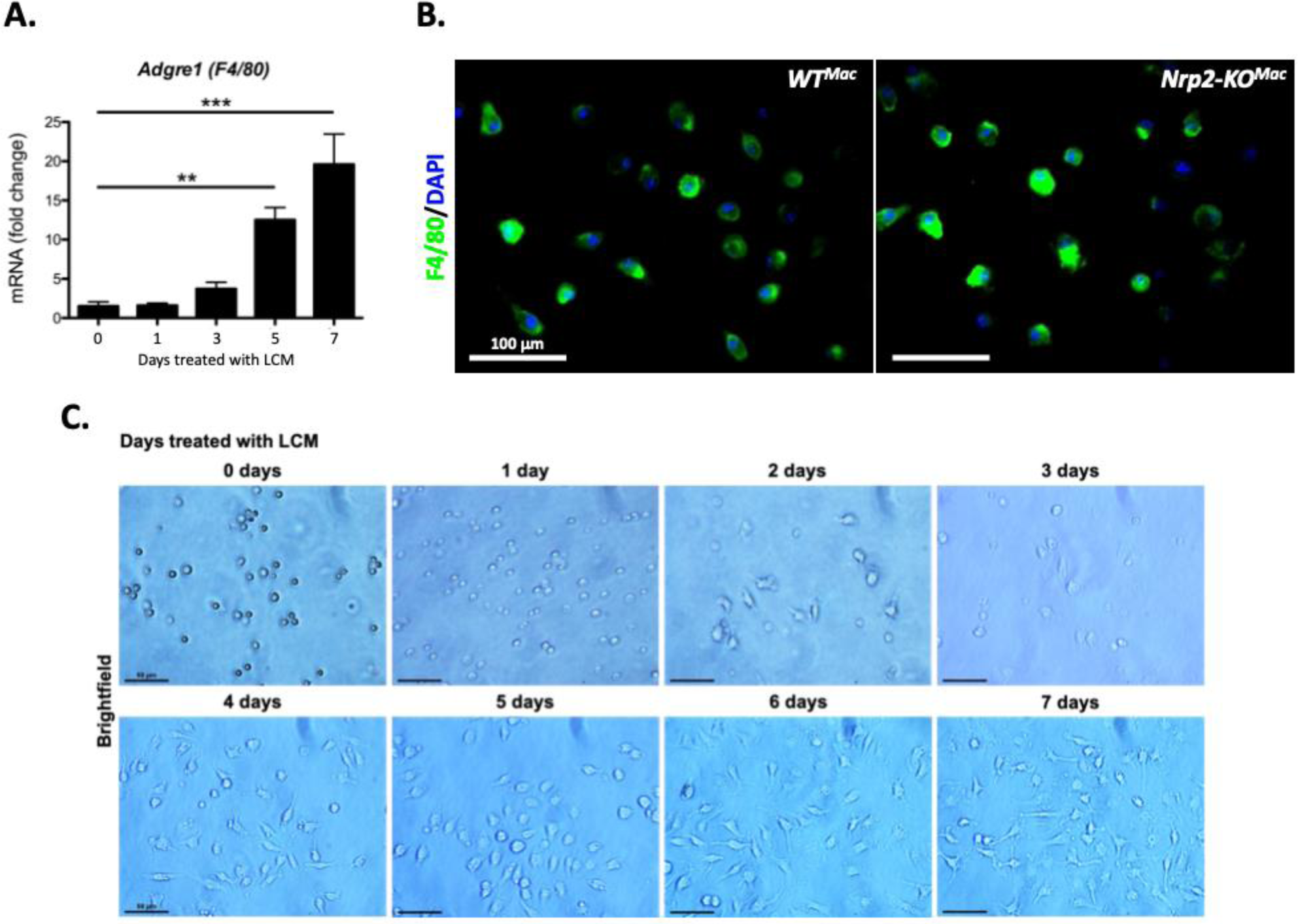
Validation of bone marrow-derived macrophage differentiation using L929-conditioned medium. **A.** Relative RT-qPCR analysis of *Adgre* (*F4/80)* expression in BMDMs after 0, 1, 3, 5 and 7 days of culture in 30% L929-conditioned medium (LCM). *Adgre* (*F4/80)* expression was normalised against expression of *β-actin* and data is represented as mean fold-change relative to expression on day 0, with error bars showing S.E.M. (n=4), ** p<0.01, *** p<0.001 (one-way ANOVA with Bonferroni post-hoc). **B.** Representative immunofluorescence images of BMDMs taken from *WT^Mac^* (left) and *Nrp2-KO^Mac^*(right) mice stained for F4/80 (green) and the nuclear stain, DAPI (blue) on day 7 of culture in 30% LCM. Cells were imaged at 40X magnification (100 μm scale bar) (n=3). **C.** Representative brightfield images of BMDMs imaged on each consecutive day (0-7) throughout culture in 30% LCM (n=3).

**Figure S2.**
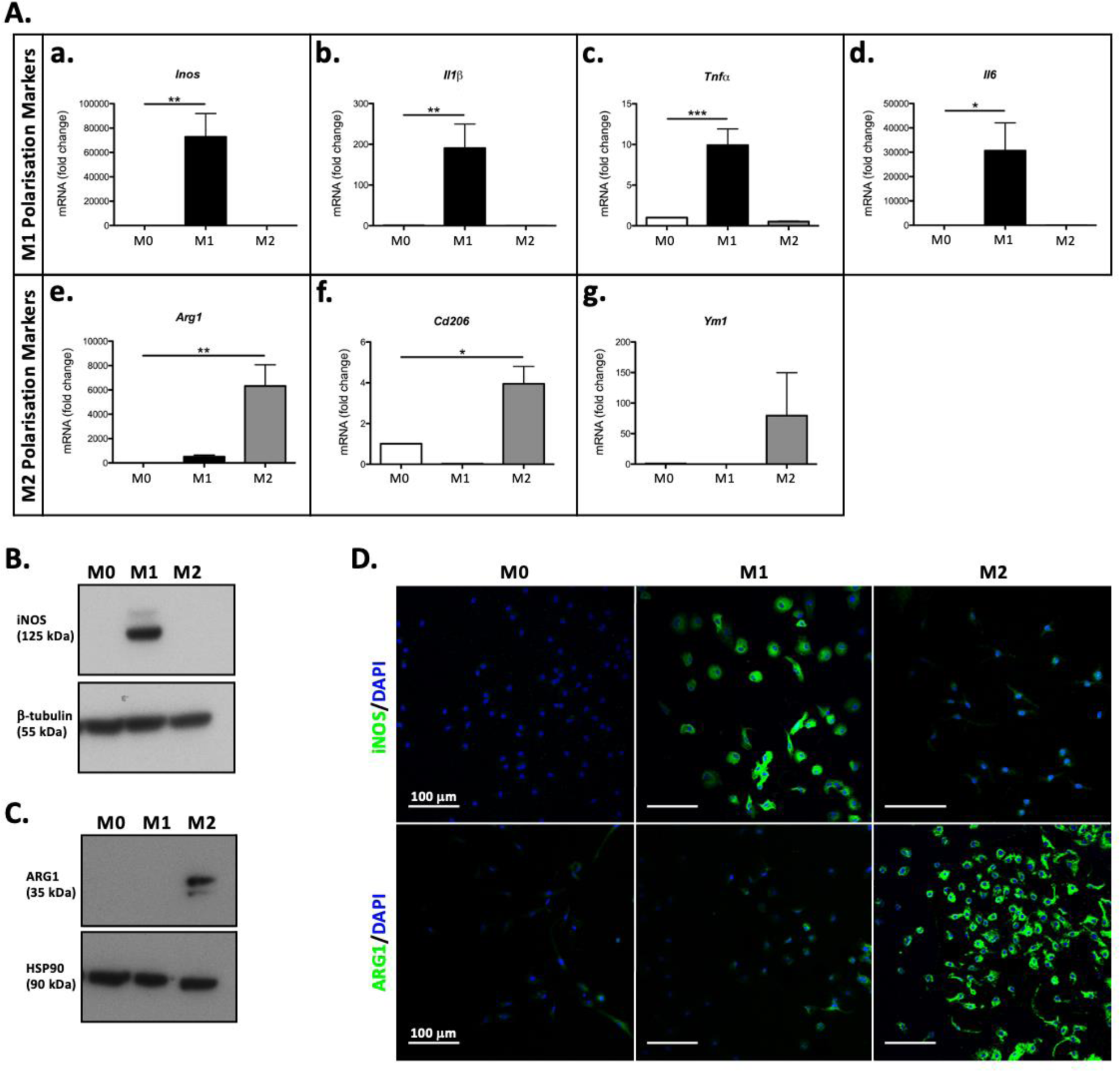
Validation of M1 and M2 polarisation in Bone marrow-derived macrophages. **A.** Relative RT-qPCR analysis of *Inos* (**a.**), *Il1β* (**b.**), *Tnfα* (**c.**), *Il6* (**d.**), *Arg1* (**e.**), *Cd206* (**f.**), and *Ym1* (**g.**) expression in M0 (serum only), M1 (LPS and IFNγ) and M2 (IL4) polarised macrophages after 24 hours of polarisation. Gene expression was normalised against *β-actin* expression and data is represented as mean fold-change relative to expression in M0 cells, with error bars showing S.E.M (n=3-5). **B-C.** Representative Western Blots showing iNOS (125 kDa) (**B.**) and ARG1 (**C.**) expression in M0, M1 and M2 polarised macrophages after 48 hours of polarisation. iNOS and ARG1 expression were normalised against β-tubulin (55 kDa) and HSP90 (90 kDa) expression, respectively (n=3). **D.** Representative immunofluorescence images of M0, M1 and M2 polarised macrophages stained for iNOS (green, top panel), ARG1 (green, bottom panel) and the nuclear stain, DAPI (blue, both panels) after 48 hours of polarisation (n=3). * p<0.05, ** p<0.01, *** p<0.001 (one-way ANOVA with Bonferroni post-hoc).

**Figure S3.**
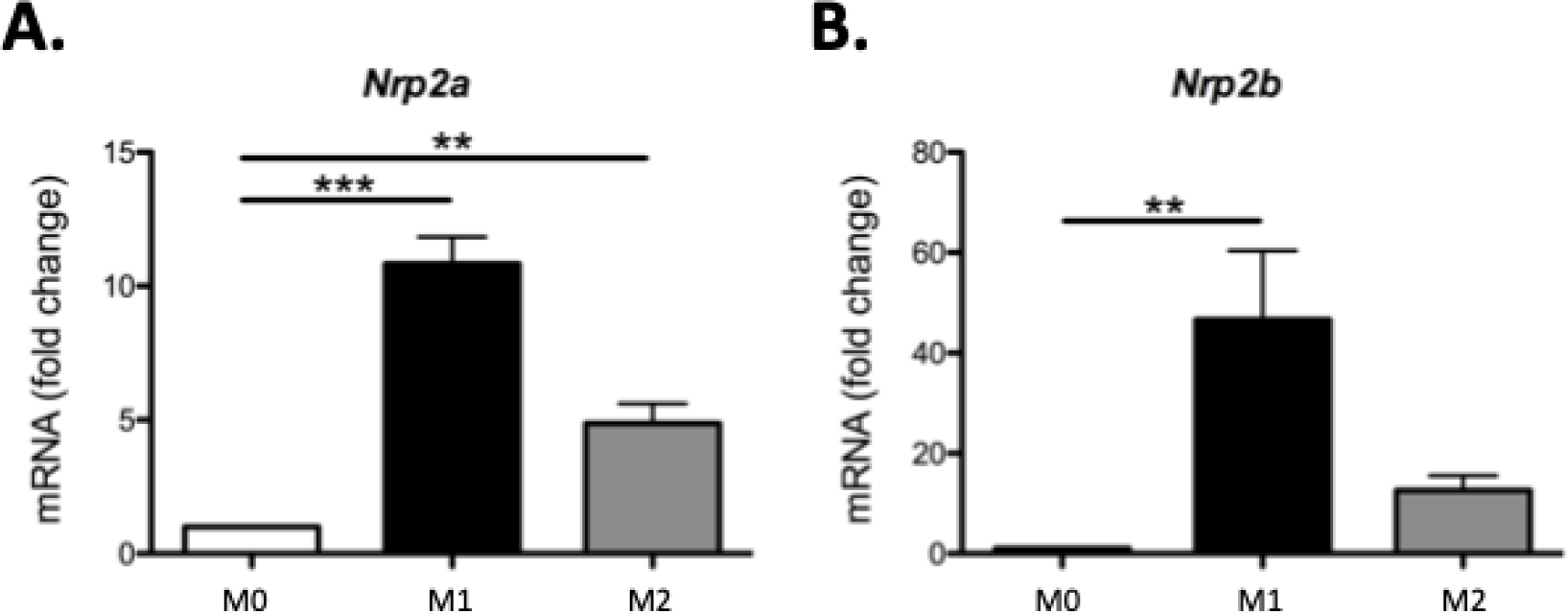
Expression of NRP2 isoforms in M1 and M2 polarised macrophages. Relative RT-qPCR analysis of *Nrp2a* (**A.**) and *Nrp2b* (**B.**) expression in M0 (serum only), M1 (LPS and IFNγ) and M2 (IL4) polarised macrophages after 24 hours of polarisation. Gene expression was normalised against *β-actin* expression and data is represented as mean fold-change relative to expression in M0 cells, with error bars showing S.E.M (n=4). ** p<0.01, *** p<0.001 (one-way ANOVA with Bonferroni post-hoc).

**Figure S4.**
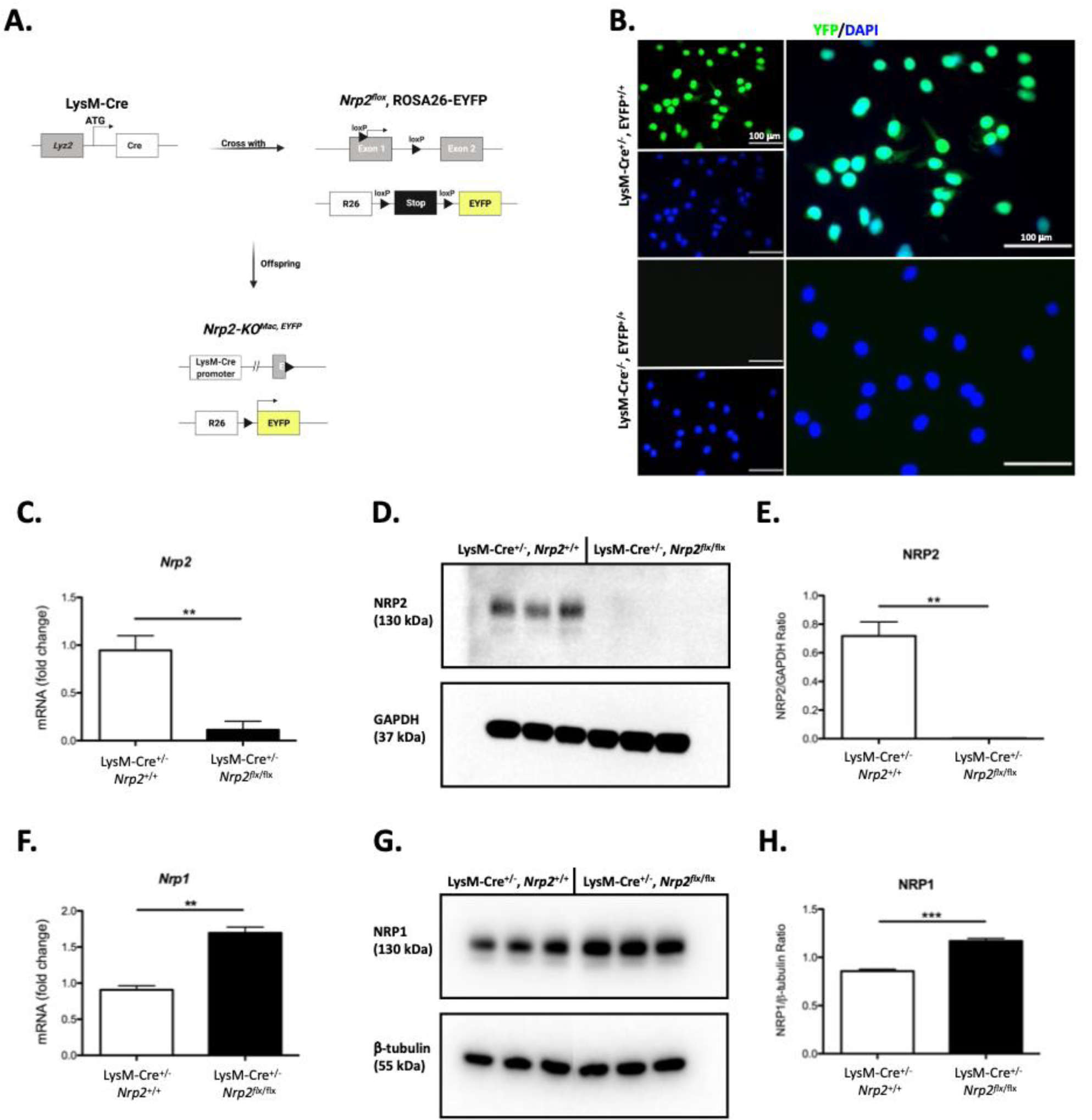
Generation of Myeloid lineage-specific Neuropilin 2 knockout mice. **A.** Mice engineered to express Cre recombinase under the control of the *Lyz2* promoter (LysM-Cre) were crossed with mice containing loxP recombination sites inserted into the *Nrp2* (within and downstream of exon 1) gene locus, and EYFP, with a loxP-flanked stop codon, within the ROSA26 (R26) locus (*Nrp2flox*, ROSA26-EYFP). Cre recombinase-mediated recombination occurs at loxP sites within the offspring, in cells expressing *Lyz2* (predominantly macrophages), resulting in excision of the loxP-flanked region and gene deletion. Excision of the loxP-flanked stop codon results in expression of EYFP. **B.** Representative fluorescence images of differentiated BMDMs extracted from LysM-Cre+/-, EYFP+/+ (top panel) and LysM-Cre-/-, EYFP+/+ (bottom panel) mice, taken on day 7 of culture in 30% LCM. Cells were stained for the nuclear stain, DAPI (blue) and YFP expression is shown in green. Merged overlays of DAPI staining and YFP expression are shown in the larger images. **C.** Relative RT-qPCR analysis of *Nrp2* expression in *WT^Mac, EYFP^* and *Nrp2-KOMac, EYFP* BMDMs. Gene expression was normalised against *β-actin* expression and data are represented as mean fold-changes relative to expression in *WT^Mac, EYFP^* cells, with error bars showing S.E.M (p=0.0096, n=3). **D-E.** Representative western blot (**D**) and quantification graph (**E**) showing NRP2 (130 kDa) expression in *WT^Mac, EYFP^* and *Nrp2-KOMac, EYFP* BMDMs. Each lane shows NRP2 expression in BMDMs extracted from an individual mouse (3 mice per genotype shown). NRP2 expression was normalised against GAPDH (37 kDa) expression and data are represented as mean NRP2/GAPDH ratios, with error bars showing S.E.M (p=0.0019, n=3). F. Relative RT-qPCR analysis of *Nrp1* expression in *WT^Mac, EYFP^* and *Nrp2-KOMac, EYFP* BMDMs. Gene expression was normalised against *β-actin* expression and data are represented as mean fold-changes relative to expression in *WT^Mac, EYFP^* cells, with error bars showing S.E.M (p=0.0013, n=3). **G-H.** Representative western blot (**G**) and quantification graph (**H**) showing NRP1 (130 kDa) expression in *WT^Mac, EYFP^* and *Nrp2-KOMac, EYFP* BMDMs. Each lane shows NRP1 expression in BMDMs extracted from an individual mouse (3 mice per genotype shown). NRP1 expression was normalised against β-tubulin (55 kDa) expression and data are represented as mean NRP1/β-tubulin ratios, with error bars showing S.E.M (p=0.0006, n=3). ** p<0.01, *** p<0.001 (unpaired, two-tailed t-test).

**Figure S5.**
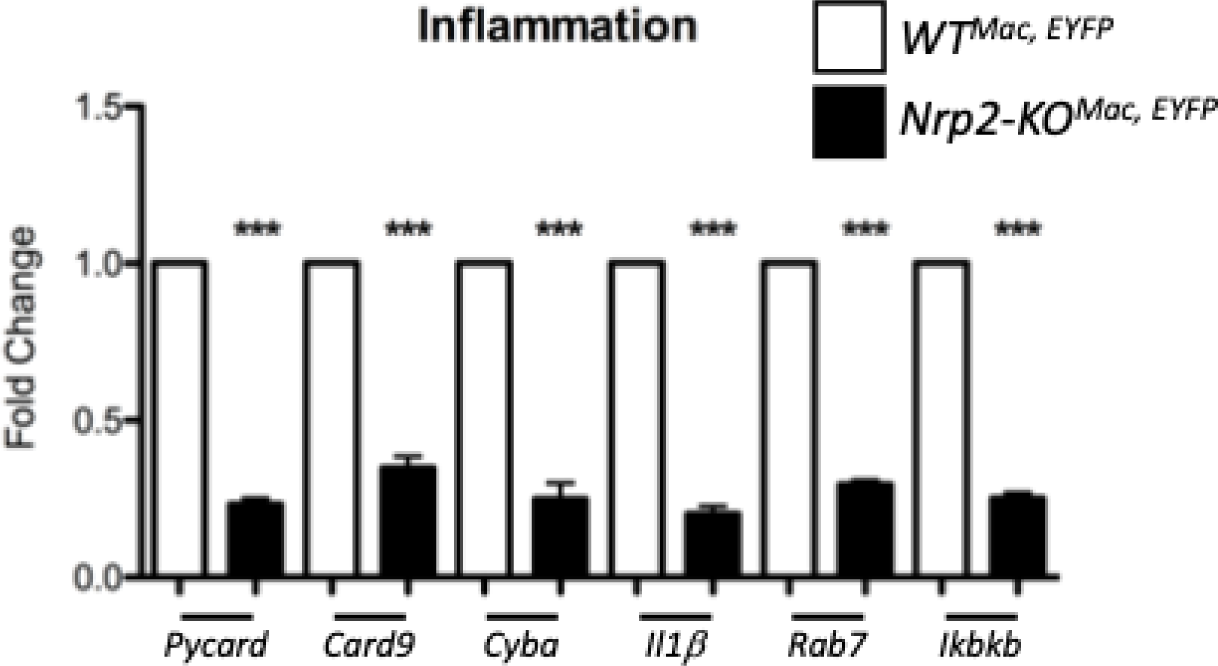
Validation of inflammatory differentially expressed genes in Neuropilin 2 knockout macrophages through RT-qPCR analysis. Relative RT-qPCR analysis of inflammation pathway gene expression in BMDMs extracted from *WT^Mac, EYFP^* (white bars) and *Nrp2-KO^Mac, EYFP^* (black bars) mice. Gene expression was normalised against expression of *β-actin* expression and data for each gene is represented as mean fold-change relative to expression in *WT^Mac, EYFP^* controls, with error bars showing S.E.M (n=3). *** p<0.001 (unpaired two-tailed t-test).

**Figure S6.**
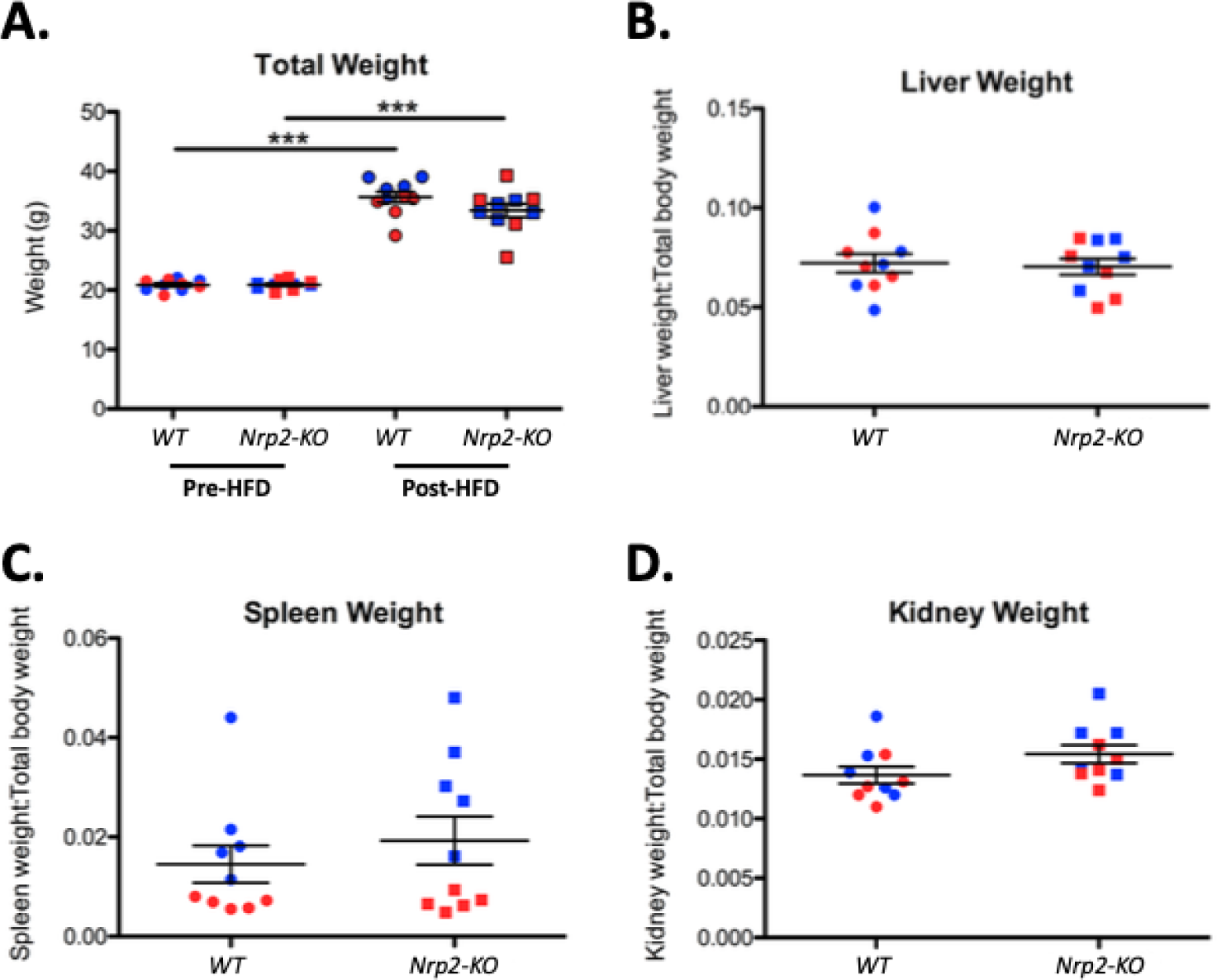
Impact of myeloid-specific Neuropilin 2 knockout on high-fat diet-induced weight gain, organ weight. **A.** Graph showing total body weight (g) of *WT^Mac, Apoe-/-, EYFP^* (*WT*, circles) and *Nrp2-KO^Mac, Apoe-/-, EYFP^* (*Nrp2-KO*, squares) before (pre-HFD) and after (post-HFD) 16 weeks of HFD feeding. Graph shows the mean +/- S.E.M (n=10) with male (blue) and female (red) mice indicated. **B-D.** Graphs showing the ratio of liver (**B**), spleen (**C**) and kidney (**D**) weight, relative to total body weight, in *WT^Mac, Apoe-/-, EYFP^* (*WT*, circles) and *Nrp2-KO^Mac, Apoe-/-, EYFP^* (*Nrp2-KO*, squares) mice after 16 weeks of HFD feeding. Graphs show mean +/- S.E.M (n=10) with male (blue) and female (red) mice indicated. *** p<0.001 (paired (**A**) and unpaired (**B-D**) two-tailed t-test).

**Figure S7:**
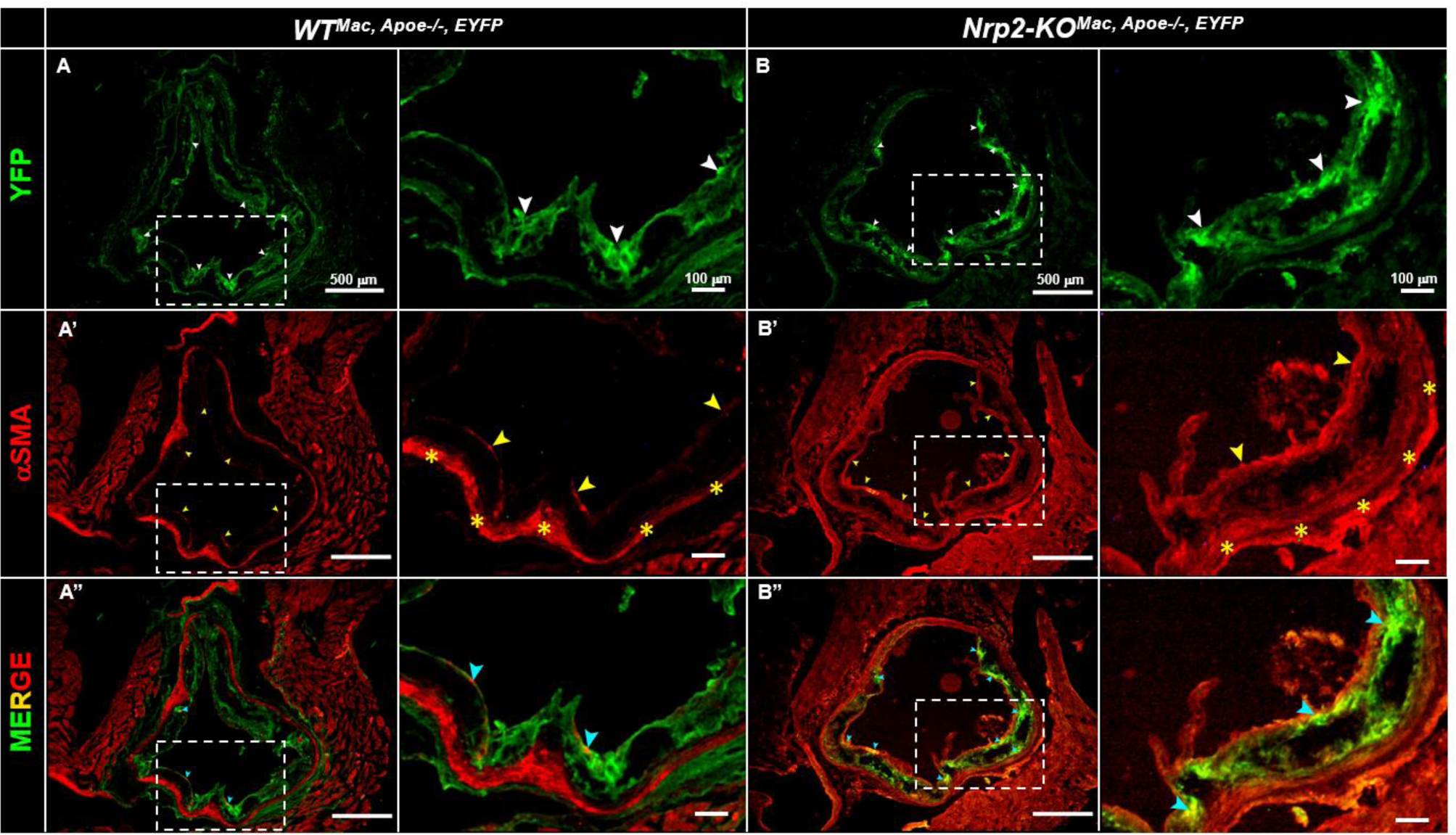
Impact of myeloid-specific Neuropilin 2 knockout on HFD-induced plaque macrophage and smooth muscle cell content. Representative images of immunofluorescence staining in the aortic roots of *WT^Mac, Apoe-/-, EYFP^* (**A-A’’**) and *NRP2- KO^Mac, Apoe-/-, EYFP^* (**B-B’’**) mice after 16 weeks of HFD feeding. YFP expression is shown in green (regions of high intensity indicated by white arrowheads) and was used to identify macrophages within the aortic roots (**A, B**). αSMA expression is shown in red (regions of high intensity in plaque cap and at luminal surface are indicated by yellow arrowheads, tunica media indicated by hollow yellow asterisks) and was used to identify SMCs within the aortic roots (**A’, B’**). Channels were merged to highlight regions of YFP and αSMA co-localisation (regions of co-localisation indicated by blue arrowheads) (**A’’, B’’**). Images are representative of staining across 2-3 aortic root sections per mouse (n>6) and areas of image highlighted by dotted rectangle are represented at higher magnification to the right (second and fourth columns for *WT^Mac, Apoe-/-, EYFP^* and *NRP2-KO^Mac, Apoe-/-, EYFP^* respectively).

